# BRO is a proteinaceous binding receptor for human coronavirus OC43

**DOI:** 10.64898/2026.11.18.767413

**Authors:** Jaewon Song, Young-Jun Park, M. Alejandra Tortorici, Li Zhang, Seokoh Moon, Amin Addetia, Vasisht V. Poosapati, Christine A. Tsu, Jack Brown, Cameron Stewart, Susan R. Ling, Benjamin Merz, Amandine Aka, Zhe Zhao, Ellen F. Foxman, Amanda M. Jamieson, Craig B. Wilen, David Veesler, Sanghyun Lee

## Abstract

*Embecoviruses* such as human coronavirus (HCoV) OC43 have long been viewed as “sialic acid–dependent” viruses that attach to cells through interactions with cell surface sialoglycans^1–5^. Here, we identify a specific proteinaceous receptor that harbors the sialoglycan determinant and also contributes protein contacts for viral spike (S) recognition. Using an unbiased surfaceome CRISPR activation screen for OC43 spike binding, we discovered that the previously uncharacterized human gene *C1ORF54* encodes this receptor, which we name *B*inding *R*eceptor for *O*C43 (BRO). BRO engages the OC43 S N-terminal domain through *O*-linked 9-*O*-acetylated sialoside determinants and neighboring protein contacts. BRO expression in human cells promotes OC43 infection, whereas BRO knockout or blockade with the BRO ectodomain or anti-BRO antibody inhibit infection, including in differentiated primary human airway cultures. BRO orthologs from diverse mammals support OC43 infection when expressed in human cells, suggesting potential contributions to OC43 zoonotic circulation and cross-species transmission. Our findings reshape our understanding of OC43 entry, and pave the way to develop countermeasures targeting this receptor interaction.

## Main text

Coronaviruses are a large group of enveloped positive strand RNA viruses infecting mammals and birds and are divided into four genera: alpha, beta, gamma, and delta^6^. Several coronaviruses are known to infect humans, such as the four endemic HCoVs, alpha-CoVs 229E and NL63, and beta-CoVs HKU1 and OC43. They typically cause mild respiratory infections but can lead to more severe disease, including bronchitis and pneumonia, especially in infants, the elderly, and immunocompromised patients^7,8^. Furthermore, three beta-CoVs have spilled over to humans since 2000 and caused outbreaks of severe and fatal respiratory syndromes^9–12^.

Coronavirus entry into cells is mediated by the trimeric spike (S) glycoprotein which is anchored in the viral membrane and decorates the surface of coronaviruses. The S glycoprotein is a key determinant of host range, tissue tropism, and pathogenesis and it is the target of neutralizing antibodies^13,14^. Receptor recognition is mediated by the S_1_ subunit, whereas the S_2_ subunit contains the fusion machinery^15,16^. The S_1_ subunit can be further divided into domain A (also called the N-terminal domain, NTD), domain B (also named receptor-binding domain, RBD), and domains C and D^14,17^. OC43, HKU1, and several other embecoviruses have a conserved glycan binding pocket within the NTD that is specific for 9-*O*-acetylated sialosides (9-*O*-Ac-Sia)^1–3^ and promotes host cell attachment as part of the entry process^4,5^. Recent findings suggested a two-step entry process for HKU1, where sialoside binding to the NTD induces S conformational changes, thereby exposing the receptor-binding motif within the RBD for engagement of its proteinaceous receptor, TMPRSS2^18–22^. A similar logic could apply to OC43, although prior structural work did not find evidence of free 9-*O*-Ac-Sia monosaccharide-induced S conformational changes^23^. Proteinaceous entry receptors identified for HCoVs include ACE2 (SARS-CoV, SARS-CoV-2, and NL63), DPP4 (MERS-CoV), APN (229E, CCoV-HuPn-2018, and PDCoV), and TMPRSS2 (HKU1)^18,24–29^, leaving OC43 as the only HCoV without a known protein receptor. Cellular entry of OC43 has been proposed to be mediated by the NTD-binding to 9-*O*-Ac-sialoglycans on the cell surface although the identity of host molecules that present the functional sialosides remains elusive.

In this study, we identify C1ORF54 (hereafter referred to as BRO, *B*inding *R*eceptor for *O*C43) as a proteinaceous binding receptor for OC43. BRO reframes OC43 entry as receptor-driven by presenting a functional 9-*O*-acetylated sialoglycan determinant while also contributing protein-protein contacts, establishing a composite glycan/protein recognition mode that confers specificity.

### *Surface*ome CRISPRa screen identifies a cellular receptor for HCoV OC43 spike protein

To identify cellular entry factors required for HCoV-OC43 infection, we performed a *surface*ome CRISPR activation (CRISPRa) screen, an unbiased receptor discovery platform, which previously identified LRRC15 as an inhibitory attachment factor for SARS-CoV-2^30^, and FcRN as an entry receptor for human astrovirus^31^. Human melanoma A375 cells stably expressing catalytically dead Cas9 (dCas) were transduced with the *surface*ome CRISPRa sgRNA library containing four sgRNAs targeting each of 6,011 genes encoding surface proteins that have transmembrane domains, GPI-motifs, or are located on the plasma membrane, along with 1,000 non-targeting controls (**Fig 1A**). After incubation with target proteins and staining with fluorescently labelled secondary antibodies, cells with high fluorescence were isolated by flow cytometry and subjected to sgRNA sequencing. While the negative control (hIgG1, human IgG1 isotype control bait) identified the *FCGR2C* Fc receptor gene (**Fig 1B**), several genes were specifically enriched in the cells binding to the Fc-tagged OC43 S_1_ subunit. Specifically, *C1ORF54* was the most enriched hit, along with *LRRC15* and two genes encoding sialyltransferases, *ST8SIA4* and *ST8SIA5* (**Fig 1C; Supplementary Table 1**). *C1ORF54* (*BRO*) is an uncharacterized gene predicted to encode a protein with a signal peptide and little known physiological functions^32^. A His-tagged SARS-CoV-2 S ectodomain trimer served as a positive control identifying both *LRRC15* and *ACE2*, consistent with our previous screens using Fc-tagged SARS-CoV-2 S_1_ protein^30^ (**Fig 1D**). Overall, the *surface*ome CRISPRa screen effectively identified host factors interacting with the OC43 (*BRO*, *LRRC15*) and SARS-CoV-2 (*LRRC15*, *ACE2*) S proteins or involved in the biosynthesis of receptor moieties (*ST8SIA4* and *ST8SIA5*).

**Figure 1.**
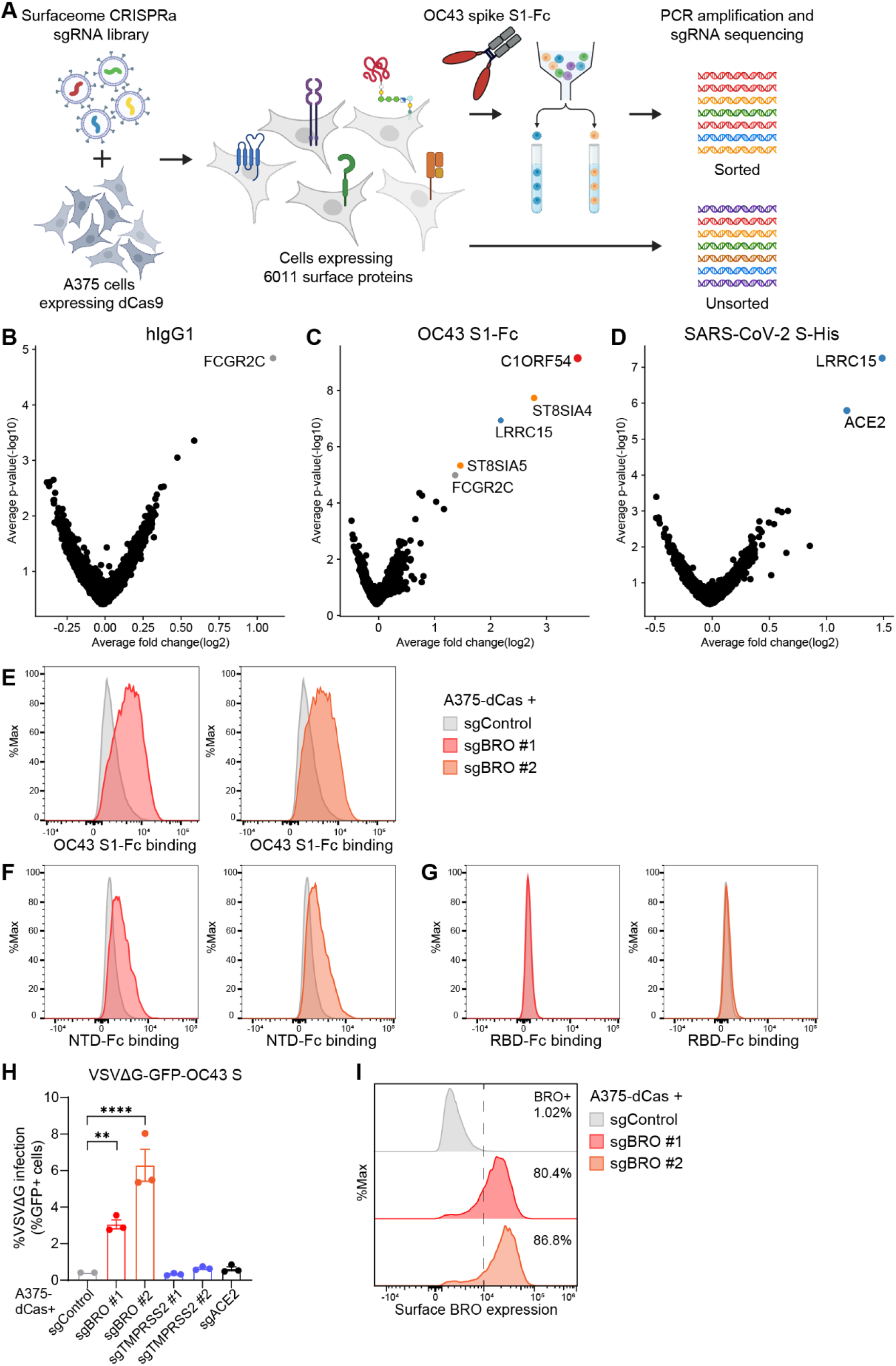
C1ORF54 (BRO) is a receptor for HCoV-OC43. (A) *Surface*ome CRISPR activation screening workflow. dCas9-expressing A375 cells were transduced with sgRNA library targeting 6,011 surface proteins, and cells binding to OC43 S_1_-hFc were sorted for sgRNA sequence analysis. (B-D) Volcano plots showing sgRNAs enriched in cells binding to human IgG control (B), OC43 S_1_-Fc (C), or the SARS-CoV-2 S trimer (D). Fold change and p-value for each gene were calculated compared to the unsorted sample. (E-G) Flow cytometry analysis of binding of OC43 wildtype S_1_-Fc (E), NTD-Fc (F), or RBD-Fc (G) to A375 cells transduced with activating sgRNAs targeting *BRO*. (H) Entry of VSVΔG-GFP particles pseudotyped with OC43 S into A375 cells transduced with the indicated activating sgRNAs. GFP signal was measured at 24 hpi by flow cytometry (n=3). Data represent means ± SEM. Data were analyzed by one-way ANOVA with Dunnett’s multiple comparisons test. \*\**p* < 0.01; \*\*\*\**p* < 0.0001. (I) A375 cells were transduced with indicated activating sgRNAs and surface BRO expression was measured by flow cytometry.

We confirmed that OC43 S_1_-Fc bound to A375 cells upon BRO induction with two independent CRISPRa sgRNAs (**Fig 1E**) and discovered that recognition was mediated by the NTD and not the RBD (**Fig 1F, G**). To understand the functional relevance of BRO recognition to the OC43 entry process, we produced vesicular stomatitis virus (VSV) particles encoding GFP and pseudotyped with the OC43 S protein (VSVΔG-GFP-OC43 S). Cell-surface expression of BRO, but not TMPRSS2 or ACE2, was sufficient to render A375 cells susceptible to OC43 pseudovirus entry (**Fig 1H, I**). Collectively, these results demonstrate that BRO is displayed at the plasma membrane, recognized by OC43 S via its NTD, and promotes OC43 S-mediated entry into cells.

### BRO is a GPI-anchored surface protein that promotes OC43 infection

Human BRO is predicted to encode a signal peptide at the N-terminus and to harbor a glycosylphosphatidylinositol (GPI)-anchoring motif with an omega-site (ω site) at residue position D107^33^ (**Fig 2A**). Following cleavage at the C-terminus of the ω+1 site, linking to the GPI moiety would be expected to tether BRO to the outer leaflet of the plasma membrane^34^. To confirm this predicted topology experimentally, we designed a dual-tagged BRO expression plasmid containing a twin Strep tag downstream of the signal peptide and a C-terminal Flag tag. Flow cytometry analysis of the BRO construct expressed in HEK293T cells enabled detection of the Strep tag both on the cell surface and intracellularly whereas the Flag tag was detected only intracellularly, indicating that the N-terminal region of BRO is displayed on the cell surface (**Fig 2B, C**).

**Figure 2.**
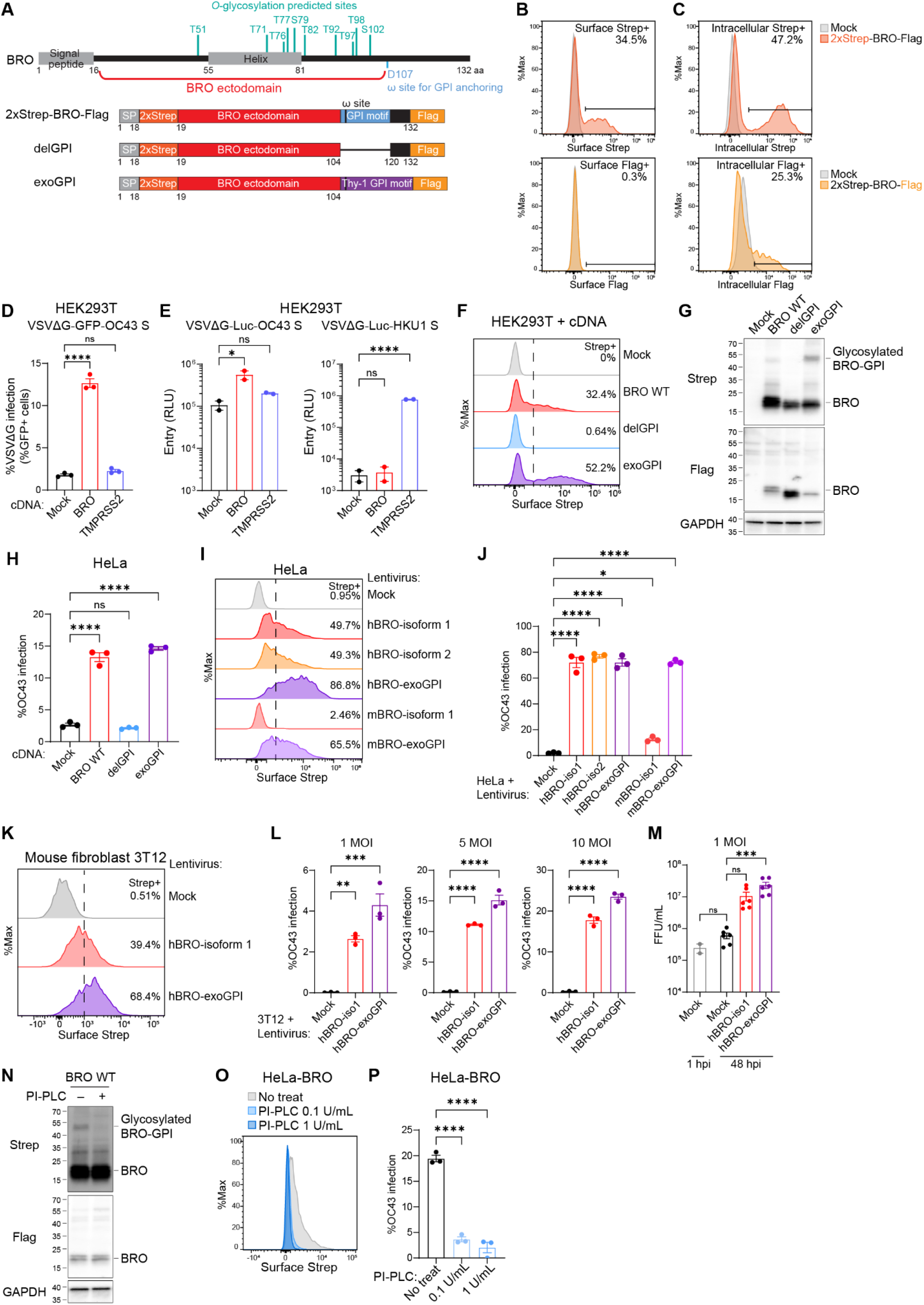
BRO is a GPl-anchored surface protein that permits OC43 infection. (A) Human BRO protein. BRO has a N-terminal signal peptide and D107 is predicted as a GPl modification site. Ten amino acids (T51, T71, T76, T77, S79, T82, T92, T97, T98, and S102) are predicted at *O*-glycosylation sites. For ectopic expression of BRO, a twin Strep tag was inserted after the signal peptide and a Flag tag was inserted at the C-terminus. (B, C) HEK293T cells were transfected with 2xStrep-BRO-Flag and Strep or Flag tag expression was measured by flow cytometry. (B) To measure surface expression, cells were stained with corresponding antibodies without permeabilization. (C) To measure intracellular expression, cells were fixed and permeabilized, then stained with antibodies. (D, E) HEK293T cells were transfected with BRO or TMPRSS2 and infected with VSVΔG-GFP (D) or VSVΔG-Luc (E) pseudotyped with OC43 or HKU1 spike. (D) Viral entry was assessed by measuring GFP signal by flow cytometry (n=3). (E) Viral entry was measured by luciferase assay, with each data point representing a biological replicate (n=2). (F, G) HEK293T cells were transfected with 2xStrep-BRO-Flag WT or to mutants, in which endogenous GPI motif is deleted (delGPI) or replaced with an exogenous GPI anchoring motif from mouse Thy-1 protein (exoGPI). (F) Expression of the Streg tag on the surface was measured by flow cytometry. (G) Expression of the Strep and Flag tag was measured by western blotting. (H) HeLa cells were transiently transfected with indicated BRO constructs and infected with OC43 virus at 5 MOl. Intracellular OC43 N expression was measured at 24 hpi by flow cytometry (n=3). (I, J) HeLa cells stably expressing indicated BRO protein with 2xStrep and Flag tag were generated. (I) Surface expression of Strep tag was measured by flow cytometry. (J) Cells were infected with OC43 virus at 1 MOI and intracellular OC43 N level was measured at 24 hpi by flow cytometry (n=3). (K-M) 3T12 cells stably expressing indicated BRO protein with 2xStrep and Flag tag were generated. (K) Surface expression of Strep tag was measured by flow cytometry. (L) Cells were infected with OC43 virus at indicated MOl and intracellular OC43 N level was measured at 24 hpi by flow cytometry (n=3). (M) Cells were infected with OC43 at 1 MOl and virus production was measured at 48 hpi by focus-forming assay (n=6 from two independent experiments). (N) HEK293T cells were transfected with 2xStrep-BRO-Flag and analyzed by western blotting after incubating with 10 U/mL of PI-PLC for 1 h. (O, P) HeLa cells stably expressing BRO were pretreated with PI-PLC at indicated concentrations for 1 h and infected with OC43 at 0.2 MOl. Surface Strep tag expression (O) and intracellular OC43 N expression (P) was measured by flow cytometry (n=3). (D, E, H, J, L, M, P) Data represent means ± SEM. Data were analyzed by one-way ANOVA with Dunnett’s multiple comparisons test. ns, not significant; \**p* < 0.05; \*\**p* < 0.01; \*\*\**p* < 0.001; \*\*\*\**p* < 0.0001.

Entry of VSV pseudotyped with OC43 S was increased in HEK293T cells by transient expression of BRO but not TMPRSS2 (**Fig 2D**). Conversely, HKU1 pseudovirus entry was detected in TMPRSS2-expressing cells but not in BRO-expressing cells (**Fig 2E**), suggesting distinct receptor usage by these closely related human coronaviruses. Enhanced OC43 pseudovirus entry and OC43 S_1_-Fc binding was also observed using HEK293T cells stably expressing BRO upon lentiviral transduction, further supporting a role for BRO in OC43 entry (**Extended Data Fig 1C-E**).

To examine the requirement for the BRO GPI anchoring motif for surface expression and viral entry, we removed the GPI anchoring motif from the expression construct (delGPI), or replaced it with an exogenous motif from a well-established GPI-anchored protein, Thy-1^35^ (exoGPI) (**Fig 2A**). Surface expression of BRO was completely abolished in delGPI-transfected cells, and restored in exoGPI cells (**Fig 2F**). Immunoblotting of these cells detected the full-length Strep-BRO-Flag at the expected molecular weight (∼20 kDa) in all three conditions. Moreover, we observed a higher molecular weight species upon transfection of BRO WT and particularly exoGPI, but not delGPI, possibly reflecting a glycosylation-mediated mobility shift of the GPI-anchored form (**Fig 2G**). GPI-anchored BRO was detected using an anti-Strep antibody, but not an anti-Flag antibody, consistent with the predicted cleavage of the C-terminus and demonstrating that BRO is a GPI-anchored cell surface protein.

We next investigated whether BRO promotes infection of authentic OC43 virus using the prototypical tissue culture-adapted strain (VR-1558; ATCC) by detecting intracellular levels of nucleocapsid (N) protein by flow cytometry. Transient ectopic expression of BRO in HeLa cells promoted OC43 infection in a GPI anchor-dependent manner (**Fig 2H**). Humans encode two isoforms of *BRO*, which are nearly identical except for a single amino acid addition to isoform 1 (isoforms 1 and 2 comprise 132 and 131 amino acids, respectively) (**Extended Data Fig 1A**). Both isoforms exhibited comparable surface expression and supported viral infection when expressed in HeLa cells. Introducing the exogenous GPI anchoring motif (exoGPI) further boosted surface expression of BRO, but provided minimal additional gain in viral entry, likely due to sufficient expression of the endogenous forms for function (**Fig 2I, J**). We confirmed that the OC43 spike S_1_ subunit and the NTD, but not the RBD, bound to HeLa cells ectopically expressing BRO (**Extended Data Fig 1F**), consistent with the CRISPRa results. We note that BRO-dependent entry was consistently lower for VSVΔG pseudovirus harboring the reference OC43 spike (VR-759) than for authentic OC43 strain VR-1558 (**Extended Data Fig 2A, B**). In a virion-binding ELISA normalized for S content, VSVΔG pseudovirus and authentic OC43 bound similarly to the OC43 NTD-directed monoclonal antibody 41F12^36^, whereas the pseudovirus showed reduced binding to soluble BRO-hFc and mucin relative to authentic virus (**Extended Data Fig 2D-G**). This difference may underlie the lower BRO-dependent entry of pseudovirus and may reflect differences in virion context and/or S sequence difference between the VR-759-based pseudovirus and the tissue culture-adapted VR-1558 virus strain^37,38^.

OC43 is phylogenetically related to rodent embecoviruses. In neonatal mice, intranasal OC43 infection can lead to lethal acute encephalitis, whereas respiratory infection in adult mice is limited and typically asymptomatic^39–41^. Human OC43 strains exhibit limited infectivity in mouse cell lines *in vitro*^42,43^. Mouse *BRO* encodes three uncharacterized isoforms, all predicted to have a GPI-anchoring motif (**Extended Data Fig 1A, B**).

Stable expression of mouse BRO isoform 1 in HeLa cells led to detectable surface expression (2.46% of cells were Strep+), albeit much lower than that of human BRO, and promoted weak OC43 infection. Replacing the mouse BRO GPI anchoring motif with the aforementioned exogenous GPI motif enhanced surface expression and OC43 infectivity, relative to wildtype mouse BRO, to levels comparable to human BRO (**Fig 2I, J**). These results indicate that the mouse BRO ectodomain plays a conserved role in supporting OC43 entry, with receptor activity limited primarily by surface presentation under these conditions.

We subsequently found that stable expression of human BRO with the wildtype or the exogenous GPI anchoring motif in mouse fibroblast 3T12 cells rendered them permissive to authentic OC43 infection and production of progeny viruses (**Fig 2K-M**). These results indicate that BRO expression is sufficient to render cells susceptible to productive OC43 infection.

To further assess the role of GPI-anchored BRO, we used phosphatidylinositol-specific phospholipase C (PI-PLC), which is an enzyme cleaving and releasing GPI-anchored proteins from the cell surface. PI-PLC treatment eliminated the glycosylated GPI-anchored form of BRO in HEK293T cells transfected with the Strep-BRO-Flag cDNA as visualized by disappearance of the species with reduced electrophoretic mobility by Western blot (**Fig 2N**). Pretreatment of HeLa cells stably expressing human BRO (HeLa-BRO) with PI-PLC reduced BRO expression on the cell surface and in turn OC43 infection (**Fig 2O, P**). Collectively, these results indicate that BRO is a GPI-anchored surface protein whose expression on the cell surface is sufficient to permit OC43 entry and replication.

### BRO is necessary for OC43 entry in human cells

To examine the role of BRO in OC43-susceptible cells, we generated clonal *BRO* knockout (KO) lines using human colon epithelial HCT-8 (also known as HRT-18) cells (**Extended Data Fig 3A**), a well-established model for OC43 infection^44^. Genome editing in the KO cells was validated by genomic DNA sequencing. OC43 infectivity was markedly reduced in two clonal *BRO* KO cell lines (clones G9 and E6) compared to the parental HCT-8 cells (**Fig 3A**) and production of progeny virus was reduced by 10- to 90-fold (**Fig 3B**). Complementation of each clone with Strep-tagged BRO rescued both OC43 infectivity and progeny production (**Fig 3A, B; Extended Data Fig 3B**). The lower BRO-dependent infectivity of the VSVΔG pseudovirus relative to authentic OC43 was also observed in HCT-8 cells, consistent with the findings made with ectopic BRO-expressing cell lines (**Extended Data Fig 2C**). The necessity of BRO for OC43 infection was also confirmed in human lung fibroblast MRC-5 cells, a widely used human cell line for OC43 studies^38^ (**Fig 3C**). Owing to challenges in isolating clonal MRC-5 cells, we generated polyclonal BRO KO populations using two different sgRNAs. ICE analysis indicated ∼56% and ∼73% editing efficiencies in each *BRO* KO population (**Extended Data Fig 3C**), leading to significantly reduced OC43 infection and production of progeny virus (**Fig 3C-E**). These results indicate that BRO is necessary for efficient OC43 infection in naturally susceptible human cells.

**Figure 3.**
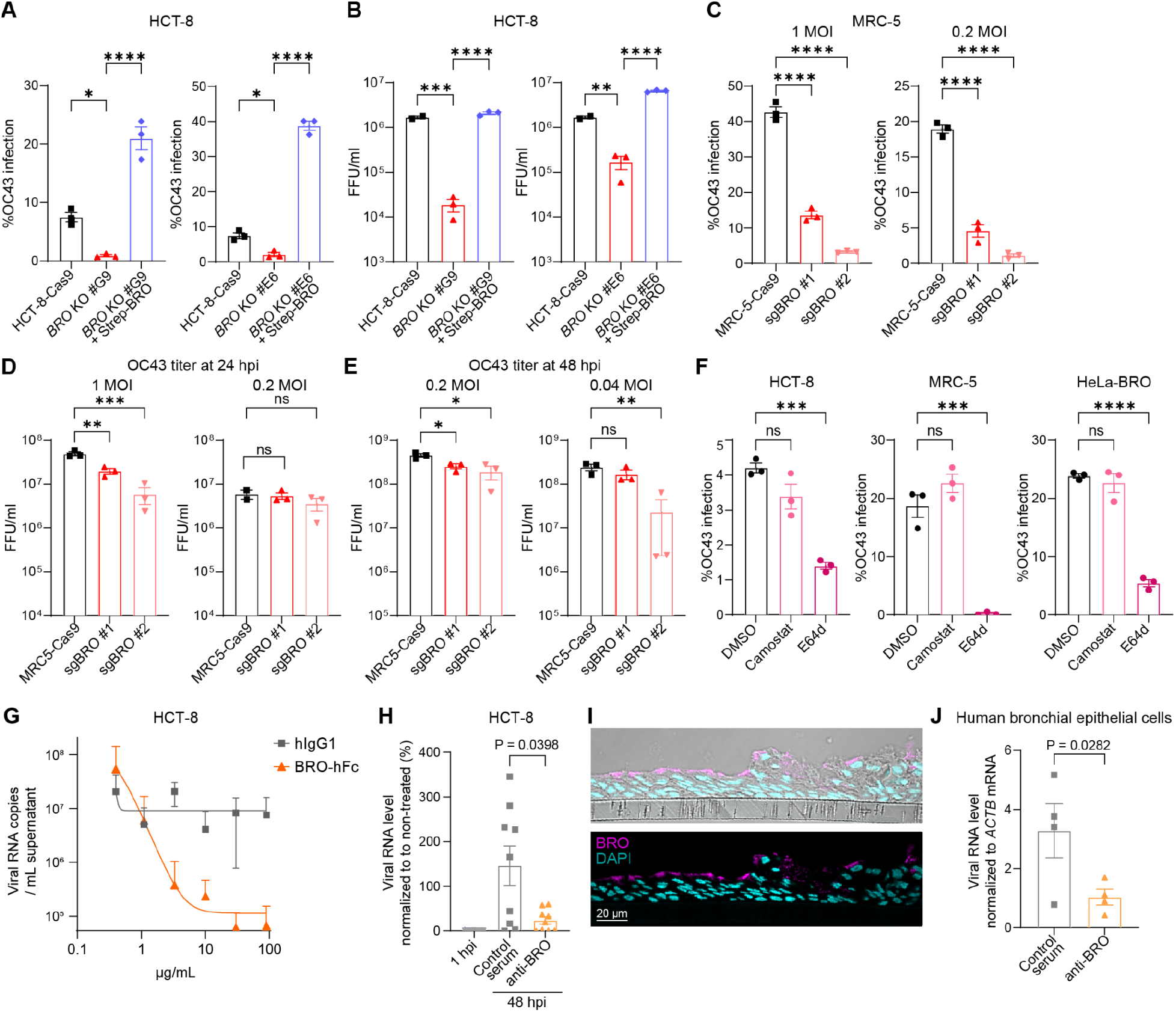
BRO is necessary for OC43 entry. (A, B) HCT-8-Cas9, *BRO* clonal KO cell lines, and KO cells complemented with Strep-BRO were infected with OC43 at 1 MOI and analyzed at 48 hpi. (A) Viral infection was measured by intracellular OC43 N staining and flow cytometry (n=3). (B) Viral production was measured by focus forming assay (n=3). (C-E) MRC-5-Cas9 cells transduced with two different sg*BRO* were infected with OC43 at indicated MOI. (C) Viral infection was measured at 24 hpi by intracellular OC43 N staining and flow cytometry analysis (n=3). (D, E) Viral production was measured at 24 hpi (B) or 48 hpi (C) by focus forming assay (n=3). (F) HCT-8, MRC-5, and HeLa-BRO stable cells were pretreated with Camostat, E64d, or vehicle for 1 h and infected with OC43. Intracellular N expression was measured at 24 hpi (MRC-5, HeLa-BRO) or 48 hpi (HCT-8) by flow cytometry (n=3). (G) HCT-8 cells were infected with OC43 (100 PFU) pre-incubated with varying concentrations of BRO-hFc (orange) or isotype control (grey). Viral RNA in the supernatant was measured by RT-qPCR at 48 hpi (n=5). (H) HCT-8 cells were pre-incubated with control or anti-BRO serum for 30 min and infected with 100 PFU of OC43. Viral RNA in the culture supernatant was measured by RT-qPCR at 48 hpi (n=9 from two independent experiments). (I) Immunofluorescence imaging of BRO (pink) and DAPI (cyan) in HBEC-ALI culture. Scale bar, 20 μm. (J) HBECs were pre-incubated with control or anti-BRO serum for 30 min and infected with OC43 at 10 MOI. Viral RNA level was measured by RT-qPCR at 48 hpi (n=4). (A-H, J) Data represent means ± SEM. Data were analyzed by one-way ANOVA with Dunnett’s multiple comparisons test (A-F) or one-sided unpaired t-test (H, J). ns, not significant; \**p* < 0.05; \*\**p* < 0.01; \*\*\**p* < 0.001; \*\*\*\**p* < 0.0001.

Coronaviruses can enter host cells through endocytosis followed by spike processing by endosomal cysteine proteases, or through TMPRSS2-mediated spike cleavage and plasma membrane fusion^45^. The OC43 VR-1558 strain used in this study has been shown to favor the late endosomal route for cell entry and does not utilize TMPRSS2^46^. Consistently, OC43 infection in HCT-8, MRC-5, and HeLa-BRO cells was specifically inhibited by E64d (cysteine protease inhibitor) but not by camostat (serine protease inhibitor), indicating that viral entry in these cells occurs through the endosomal pathway in our assays (**Fig 3F**).

We next set out to target BRO to inhibit OC43 infection. First, the recombinant BRO ectodomain fused to an Fc-tag (BRO-hFc) was pre-incubated with authentic OC43 prior to infection of HCT-8 cells, and viral RNA replication was quantified by RT-qPCR. OC43 pre-incubation with BRO-hFc decreased replication in HCT-8 cells in a concentration-dependent manner, suggesting that BRO-hFc acts as a decoy receptor (**Fig 3G**). Second, HCT-8 cell pre-treatment with anti-BRO serum prior to OC43 infection significantly reduced viral infectivity (**Fig 3H**). To assess the role of BRO in a physiologically relevant *ex vivo* human tissue model, we employed primary human bronchial epithelial cells (HBECs) differentiated at the air-liquid interface (ALI). Confocal immunostaining demonstrated apical expression of BRO in airway epithelial cells (**Fig 3I**). Differentiated HBECs were treated with anti-BRO serum prior to the apical inoculation of OC43 and viral replication was measured by RT-qPCR after 48 hours. We found that OC43 infection was significantly reduced with anti-BRO serum treatment (**Fig 3J**). Collectively, these results demonstrate that BRO is necessary for OC43 entry in human cells *in vitro* and in primary human airway epithelium, and that BRO is a target for receptor blockade.

### Identification of key peptide motifs and *O*-linked glycosylation sites in BRO mediating OC43 entry

Given that OC43 utilizes its NTD to engage 9-*O*-Ac-sialoside receptors and BRO, we hypothesized that BRO may harbor the sialoside moiety recognized for viral entry. The BRO ectodomain harbors ten predicted *O*-glycosylation sites, including position T51 which was previously identified in an *O*-glycoproteome study^47,48^ (**Fig 4A**) and no predicted *N*-linked glycosylation sites. We mutated individual predicted *O*-glycan sites (via serine/threonine to alanine substitutions) and quantified OC43 infection in HeLa cells transfected with each mutant. Surface expression of BRO was generally reduced by mutating single or multiple glycosylation sites, whereas intracellular levels remained largely comparable to WT BRO except for a few select mutants (**Extended Data Fig 4A-D**). Mutation of individual glycosylation sites decreased OC43 infection to similar levels across mutants (**Fig 4B**). Substitution to alanine of putative *O*-linked glycosylation site residues that are closely clustered in sequence (5mut and 4mut) led to similar entry reductions to single mutants whereas mutation of all ten sites (10mut) dampened OC43 infectivity to a greater extent (**Fig 4B**). We next rescued individual glycan sites by reintroducing the wildtype serine or threonine residues in the 10mut background. Restoring T51 in the 10mut construct significantly enhanced OC43 infectivity relative to the parent construct whereas rescuing other single glycan sites had minimal effects (**Fig 4C**).

**Figure 4.**
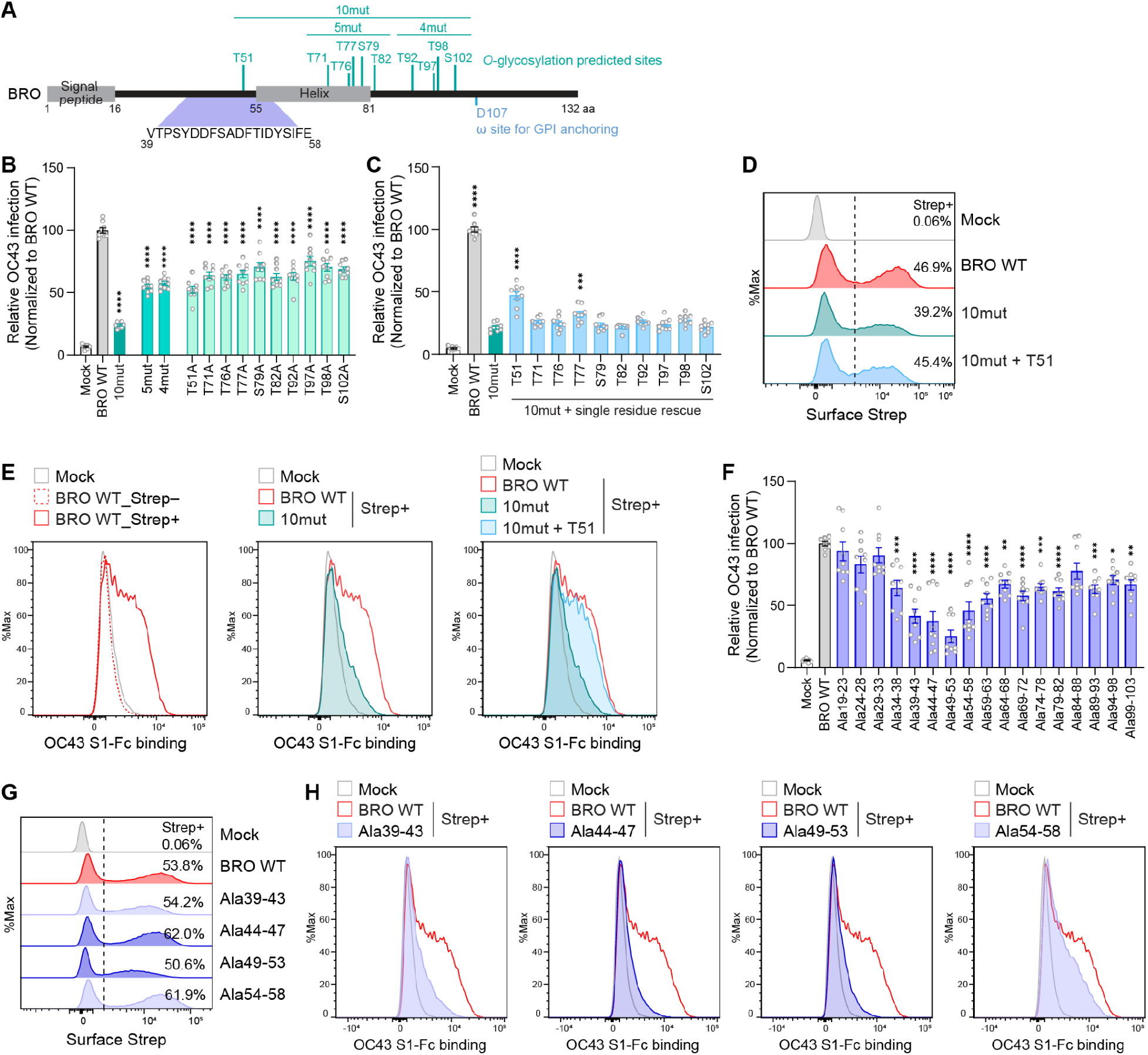
Identification of BRO molecular determinants required for OC43 entry. (A) Human BRO protein. Ten amino acids (T51, T71, T76, T77, S79, T82, T92, T97, T98, and S102) are predicted as *O*-glycosylation sites. Alanine scanning identified residues required for OC43 infection. (B, C) HeLa cells were transfected with BRO WT or indicated mutants and infected with authentic OC43 virus (VR-1558) at 1 MOI followed by intracellular N staining at 24 hpi (n=9 from three independent experiments). (B) Residues predicted as *O*-glycosylation sites were substituted with alanine. 10mut is all ten predicted glycosylation residues mutated to alanine, 5mut is T71/T76/T77/S79/T82 mutated, and 4mut is T92/T97/T98/S102 mutated. (C) Single residues of 10mut were rescued back from alanine to threonine or serine as indicated. (D, E) HEK293T cells were transfected with BRO WT or indicated mutants and incubated with OC43 spike S_1_-Fc protein, followed by flow cytometry analysis. (D) Surface Strep tag expression was measured. (E) S_1_-Fc binding to Strep– or Strep+ population was measured for cells transfected with indicated BRO WT or mutants. (F) Alanine scanning mutants were generated by mutating five consecutive residues in the ectodomain. Numbers indicate amino acid residues substituted by alanine. HeLa cells were transfected with BRO WT or indicated mutants and infected with OC43 virus at 1 MOI followed by intracellular N staining at 24 hpi (n=9 from three independent experiments). (G, H) HEK293T cells were transfected with BRO WT or indicated mutants and incubated with OC43 spike S_1_-Fc protein, followed by flow cytometry analysis. (G) Surface Strep tag expression was measured. (H) S_1_-Fc binding to Strep– or Strep+ population was measured for cells transfected with indicated BRO WT or mutants. (B, C, F) Data represent means ± SEM. Data were analyzed by one-way ANOVA with Tukey’s multiple comparisons test and p-values compared to BRO WT samples are shown. \**p* < 0.05; \*\**p* < 0.01; \*\*\**p* < 0.001; \*\*\*\**p* < 0.0001.

Because the magnitude of infection (**Fig 4B-C**) generally tracked with the level of surface BRO expression (**Extended Data Fig 4A-B**), suggesting contribution of *O*-glycosylation to BRO surface presentation, we next sought to separate effects on BRO surface expression from effects on spike binding. We investigated the role of BRO glycosylation for its receptor function by evaluating OC43 S_1_-Fc binding to cells transfected with various BRO constructs (**Fig 4D**). For this analysis, we used HEK293T cells, which showed higher and more uniform surface BRO expression across constructs, and we pre-gated on surface Strep+ cells to assess spike binding within the BRO-presenting population. OC43 S_1_-Fc bound exclusively to Strep+ cells, confirming surface BRO expression is required for OC43 S recognition (**Fig 4E**, left). In the Strep+ population, OC43 S_1_ binding was abrogated by 10mut expression (**Fig 4E**, middle) and this defect was restored by the T51-rescue construct (**Fig 4E**, right), despite slightly lower surface Strep intensity than WT. Collectively, these results suggest that *O*-glycosylation of BRO contributes to surface presentation and spike engagement, with T51 acting as a dominant determinant for S binding and viral entry.

To investigate the possible contribution of BRO peptide residues to OC43 S binding, we generated seventeen mutants, each harboring five consecutive residues substituted to alanine. We observed that mutating residues 39-58, surrounding the T51 glycan residue, led to a substantial reduction of OC43 infectivity whereas other mutants had marginal effects on OC43 infection (**Fig 4F; Extended Data Fig 4E, F**). Accordingly, cells expressing these alanine mutants exhibited reduced OC43 S_1_-Fc binding (**Fig 4G, H**), indicating that the amino acid residues (39-58) neighboring the T51 glycan site may be involved in OC43 S binding and viral entry either directly or through promoting of proper folding.

### Molecular characterization of binding between OC43 spike and BRO

To understand the molecular basis of OC43-mediated receptor engagement, we determined a single particle cryoEM structure at 2.0Å resolution of a complex between prefusion OC43 S and BRO-mFc (**Fig 5A; Extended Data Fig 5; Extended Data Table 2**). We subsequently used symmetry expansion, focused classification and local refinement to improve map resolvability around the NTD sialoside binding site, yielding a second reconstruction at 2.8Å resolution. The map resolves a di-sialoside bound to the OC43 NTD with a configuration similar to that observed in a previously described α2,8-linked 9-*O*-acetylated disialoside (9-*O*-Ac-Sia(α2,8)Sia)-bound HKU1 S structure (**Fig 5B-D**). In contrast to HKU1, however, ligand binding does not lead to opening of the RBDs, which remain in the closed state in our structure, as also observed for OC43 S bound to the monosaccharide 9-*O*-Ac-SIA^1^. Additional density corresponding to the proteinaceous part of BRO could be resolved at much lower resolution, possibly as a result of its predicted disordered nature (**Fig 5A-B**). Nevertheless, the map suggests that BRO also contacts the OC43 NTD around residues V84 and H183 which are close to each other (**Fig 5B**). Analysis of OC43 isolate sequences publicly available indicates that H183 is strictly conserved whereas position 84 most frequently harbors an aspartate residue instead of a valine (**Fig 5E**). To validate the observed binding mode, we investigated the impact on BRO binding of the OC43 NTD W90A mutation, which was previously shown to abrogate 9-*O*-Ac-SIA binding^1,2^. Disruption of the sialoside binding site through the W90A residue substitution abrogated binding of OC43 S_1_-Fc to cell surface-expressed BRO and to the BRO ectodomain (**Fig 5F-H**), consistent with the structural data. Furthermore, neuraminidase or BCoV hemagglutinin-esterase (BCoV-HE) pretreatment of BRO abolished binding to OC43 S_1_-Fc and infectivity of OC43 virus (**Fig 5G-I; Extended Data Fig 6G-H**), underscoring the key role of sialoside-mediated interactions, and particularly the 9-*O*-acetyl modification, for receptor recognition.

**Figure 5.**
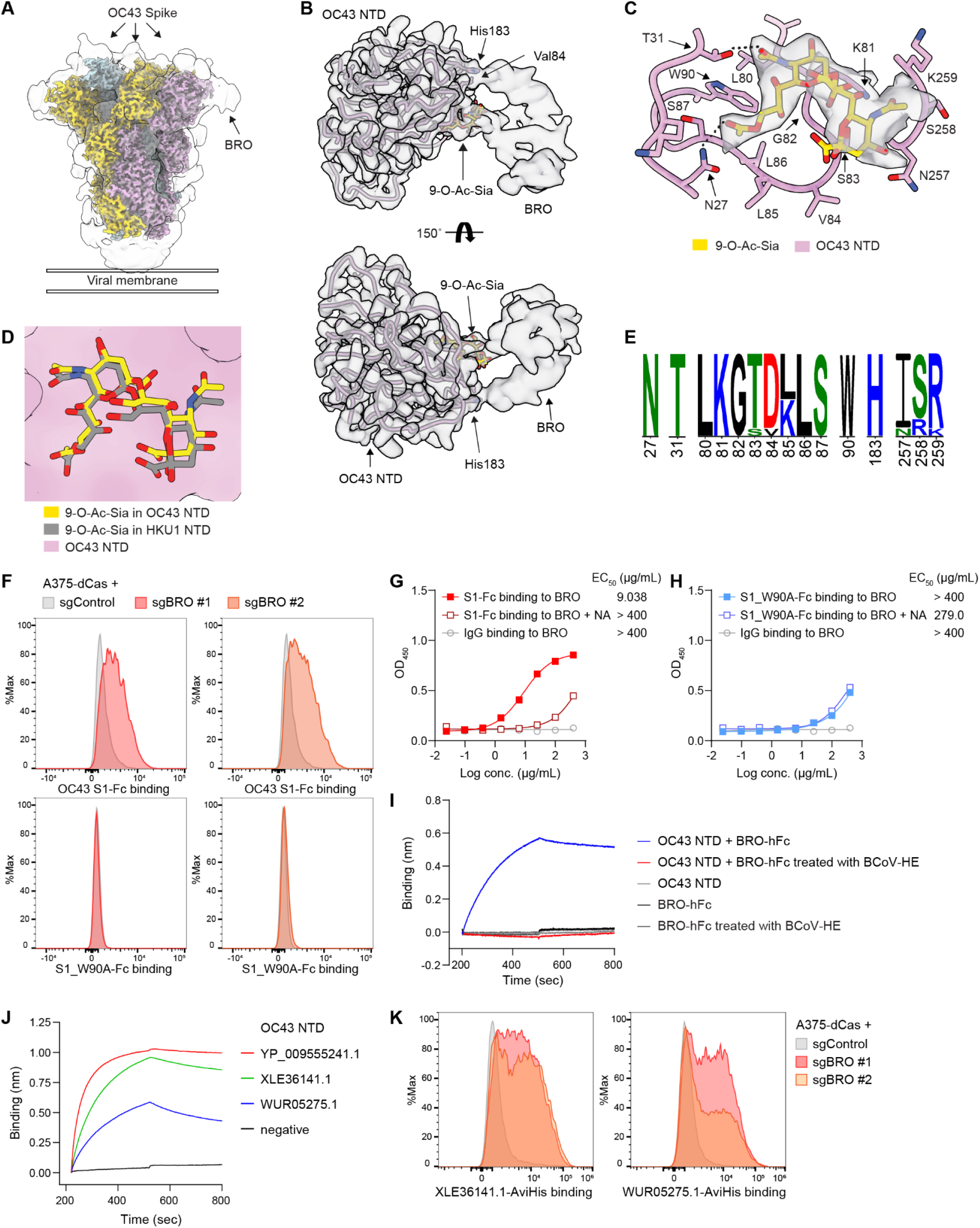
OC43 NTD directly binds to glycan on BRO protein. (A) CryoEM reconstruction of prefusion OC43 S bound to the human BRO ectodomain at 2.0 Å resolution. The sharpened map is colored by protomers. The 8Å low-pass filtered unsharpened map is shown in grey. (B) CryoEM reconstruction obtained by symmetry-expansion and local refinement of the OC43 NTD and BRO region (deconvoluted map) at 2.8 Å resolution. (C) Zoomed-in view of the modeled BRO sialoside (9-O-Ac-Sia(α2,8)Sia) bound to the OC43 NTD (deconvoluted map shown around the ligand). (D) Superimposition of the (9-O-Ac-Sia(α2,8)Sia)-bound OC43 (this study, in plum and gold) and HKU1 NTD (PDB ID:9BTB, in light blue and gray) underscoring the similar binding mode. (E) Polymorphism among 202 OC43 isolates of NTD residues interacting with 9-*O*-Ac-Sia(α2,8)Sia and the BRO proteinaceous part shown as logoplot. Amino acids are colored according to their chemical properties: hydrophobic (black), polar non-charged (green), positively charged (blue) and negatively charged (red). The residue numbering accordings to HCoV OC43 (Uniprot ID:Q696P8). (F) Flow cytometry analysis of binding of OC43 wildtype or W90A S_1_-Fc to A375 cells transduced with activating sgRNAs targeting BRO. (G, H) Binding of wildtype (G) or W90A (H) OC43 S_1_-Fc to human BRO pre-incubated or not with neuraminidase (NA) was measured by ELISA. Half-maximal effective concentrations (EC_50_) are shown. (I) Biolayer interferometry analysis of binding of BRO-hFc pre-incubated or not with BCoV HE to immobilized OC43 NTD. (J) Biolayer interferometry analysis of binding of BRO-hFc to immobilized biotinylated OC43 NTD of the prototypical strain or clinical isolates. NCBI accession numbers for the spike proteins are shown. (K) Flow cytometry analysis of binding of OC43 clinical isolate S trimers to A375 cells transduced with activating sgRNAs targeting BRO.

To investigate the possible contribution of BRO to cell surface attachment of OC43 isolates distinct from VR-1558, we assessed binding to NTDs of the OC43 clinical isolates (S protein accession numbers XLE36141.1 and WUR05275.1). We found that BRO-hFc bound to all strains tested by biolayer interferometry, with the greatest magnitude observed for the reference strain, and that the corresponding prefusion S trimers bound to cell-surface expressed BRO by flow cytometry (**Fig 5J-K; Extended Data Fig 6I**). These results suggest that BRO contributes to attachment of a broad spectrum of OC43 isolates to the surface of target cells, and in turn viral entry.

### OC43 has broad BRO mammalian species tropism

All human coronaviruses have zoonotic origins and spill over to humans through cross-species transmission, either directly or via intermediate hosts^49,50^. Embecoviruses, including OC43 and HKU1, are thought to originate from rodents^51,52^ with cattle as intermediate hosts for OC43, which diverged from a shared ancestor with bovine CoV (BCoV) around 1890^53^. Given these evolutionary relationships, we set out to understand whether BRO mammalian orthologs can promote OC43 entry.

BRO is widely conserved across mammals, including predicted signal peptide and GPI-anchoring signal (**Fig 6A**). We synthesized expression constructs of BRO orthologs from 29 mammalian species and confirmed protein expression in human cells (**Extended Data Fig 7A; Supplementary Table 2**). To investigate retention of receptor function, we assessed the ability of each BRO ortholog to promote OC43 infection when expressed on the surface of (otherwise non-permissive) HeLa cells. We found that a broad range of BRO orthologs supported OC43 entry with human (*H. sapiens*) BRO exhibiting the highest efficiency (**Fig 6A**). Mouse (*M. musculus*), rat (*R. norvegicus*) and bovine (*B. taurus*) BRO orthologs also supported robust OC43 infection, consistent with their proposed links to OC43 emergence. Among the 29 species tested, three BRO orthologs (*C. dromedarius*, *O. cuniculus*, *E. caballus*) lack a predicted signal peptide and four orthologs (*E. caballus*, *V. vulpes*, *C. lupus*, *N. procyonoides*) lack a GPI anchoring motif. These orthologs were defective in permitting OC43 infection (**Fig 6A**). Pig (*S. scrofa*) and common shrew (*S. araneus*) BRO also showed limited OC43 infection, likely due to inefficient surface expression, consistent with a low-confident prediction for signal peptide (pig) or low expression (shrew) in HeLa cells (**Fig 6A; Extended Data Fig 7B**). Most of the *O*-glycosylation sites predicted with human BRO are conserved in these mammalian species and the key residue T51 is conserved in all 29 species (**Fig 6B**). Together, these results indicate that BRO may function as a broadly conserved receptor for OC43 and suggest that it may have contributed to OC43 zoonotic circulation and cross-species transmission.

**Figure 6.**
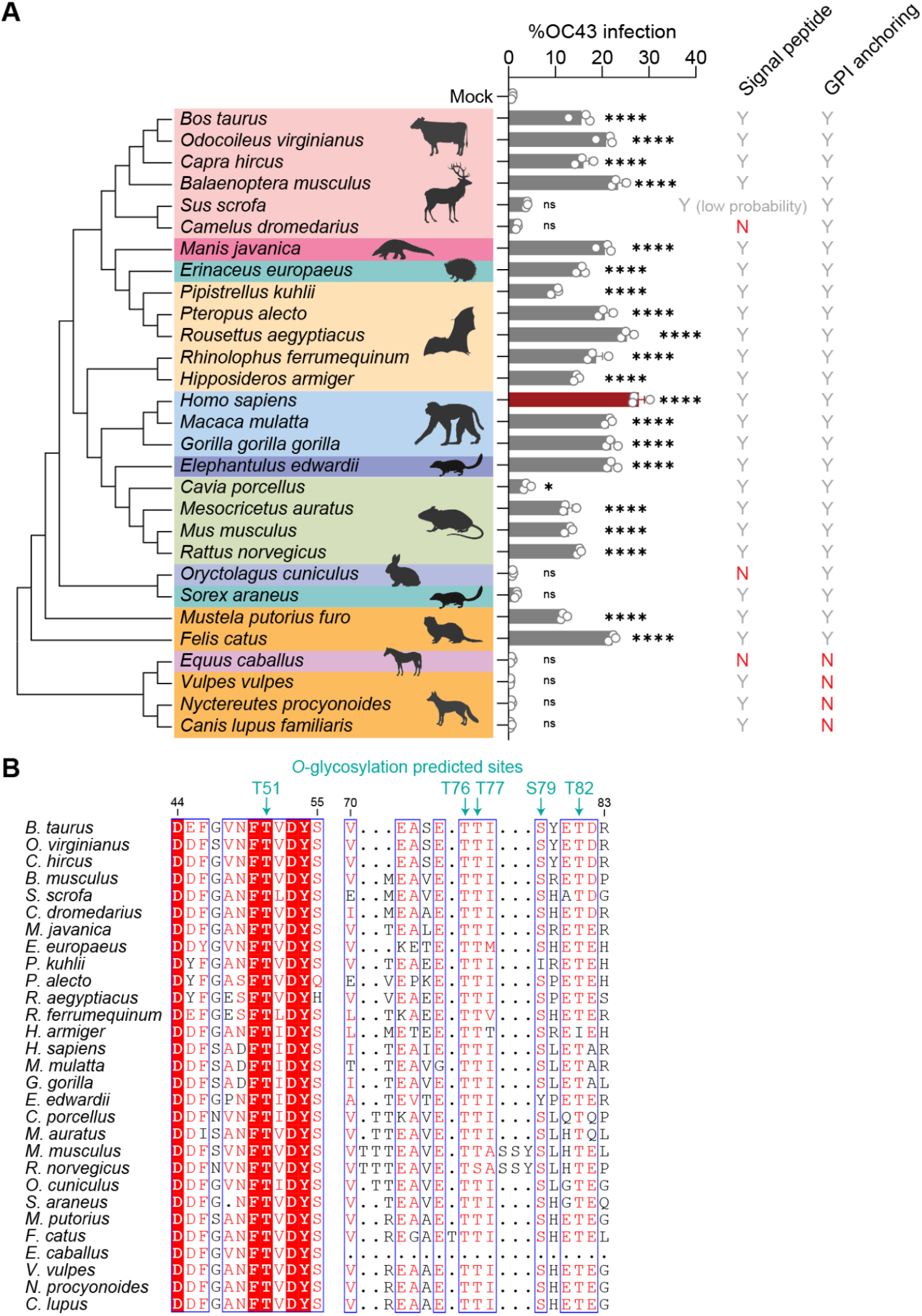
BRO from different mammalian species permit OC43 infection. (A) (Left) Phylogenetic analysis of mammalian BRO orthologs based on amino acid sequences with Orders indicated. (Middle) HeLa cells were transfected with indicated BRO ortholog constructs and infected with OC43 virus at 1 MOI followed by intracellular N staining at 24 hpi (n=3). One representative result out of two independent experiments is shown. (Right) Signal peptides and GPI anchoring was predicted for each ortholog using SignalP 6.0 and NetGPI 1.1 and shown on the right. Y, predicted; N, not predicted. Data represent means ± SEM. Data were analyzed by one-way ANOVA with Dunnett’s multiple comparisons test and p-values compared to mock transfection sample are shown. ns, not significant; \**p* < 0.05; \*\*\*\**p* < 0.0001. (B) Amino acid residues in BRO orthologs at *O*-glycosylation sites predicted with human BRO.

## Discussion

In this study, we identified BRO as a previously unrecognized, binding receptor for HCoV-OC43 using an unbiased *surface*ome CRISPRa “receptor hunting” platform. BRO promotes OC43 entry and productive infection in human cells, including human airway epithelial cultures, and BRO-targeted interventions using soluble BRO or anti-BRO antibodies inhibit HCoV-OC43 in a concentration-dependent manner.

Although OC43 has long been considered “sialic acid–dependent,” this has typically been interpreted as recognition of a defined 9-*O*-acetylated sialoglycan (including disialosides), while the identity and functional requirement of the molecular scaffold presenting the glycan has remained unknown. The discovery of BRO reframes this view by demonstrating that OC43 can exploit a defined host gene product that presents the appropriate sialylated determinant and also contributes additional receptor features beyond glycan display. Therefore, BRO represents a mechanistic bridge between purely glycan-mediated attachment and canonical receptor-driven entry.

BRO usage distinguishes OC43 from the closely related embecovirus HKU1. BRO promoted entry mediated by OC43 S but not HKU1 S despite the conserved engagement of 9-*O*-acetylated sialosides by the two viruses, possibly due to the specific contacts made by BRO amino acid residues and the HCoV-OC43 NTD. Moreover, HKU1 utilizes TMPRSS2 as an entry receptor whereas no additional receptor has been identified to bind to the HCoV-OC43 domain B. These data argue against a shared glycan receptor logic across embecoviruses despite their conserved NTD sialic acid binding pockets, and instead suggest that OC43 and HKU1 have diverged in how they couple sialic acid recognition to downstream entry steps.

A central conceptual advance from our work is that BRO functions as a sialoglycan-bearing protein receptor rather than simply increasing the bulk abundance of permissive glycans on the cell surface. Multiple observations support this conclusion. First, OC43 NTD binding to BRO required an intact NTD glycan-binding pocket (as shown with the W90A disruption) and a 9-*O*-acetyl-sialoside receptor (as demonstrated by HE-treatment of BRO), demonstrating that receptor engagement is fundamentally linked to sialic acid recognition. Second, BRO is GPI-anchored, positioning its ectodomain for presentation at the outer leaflet of the plasma membrane where S interactions with sialosides occur. We note that the recently discovered porcine haemagglutinating encephalomyelitis virus entry receptor (DPEP1) and the bat CcCoV-KY43 entry receptor (CEACAM6) are also GPI-anchored proteins^54,55^. Third, *O*-glycosylation landscape of BRO is functionally important: alanine substitution across predicted *O*-glycosylation sites reduced OC43 infection, and rescue experiments pinpointed T51 as a predominant contributor capable of restoring binding and entry in an otherwise glycan-deficient background. Lastly, cryo-EM analysis suggests that a conserved histidine motif in OC43 S mediates protein-protein interaction with BRO. Collectively, these results indicate that OC43 does not merely require any 9-*O*-Ac-Sia on the plasma membrane, but rather benefits from a structured receptor scaffold that presents an *O*-glycan determinant in an appropriate local context. The identification of sialyltransferases among the CRISPRa-enriched hits further supports the notion that glycan context shapes receptor function. While BRO can be sufficient in multiple cell types, the precise composition and modification of *O*-glycans (including 9-*O*-acetyl sialylation) likely modulates binding avidity and infection efficiency. Thus, OC43 entry appears to be governed by an interplay between a specific host receptor protein and the broader glycosylation machinery that decorates it.

Endemic human coronaviruses are ultimately zoonotic in origin. Embecoviruses are thought to have circulated in non-human hosts prior to emergence in humans. We observed that BRO orthologs from a wide range of mammals can support OC43 infection when expressed in human cells. This suggests that BRO-like proteins could function as a broadly utilized receptor platform, potentially lowering species barriers. The requirement for a signal peptide and GPI-anchoring motif emphasizes that cell-surface presentation is essential for receptor activity as orthologs lacking these features were non-functional. Conservation of predicted *O*-glycosylation features, including the key T51 position, further supports the idea that BRO provides an evolutionarily conserved scaffold for presenting permissive sialoglycans. Moreover, differences in surface expression (as observed for murine BRO and its functional improvement by the exogenous GPI motif replacement) suggest that quantitative or qualitative variation in receptor presentation could modulate susceptibility across species or tissues. In natural settings, BRO-driven permissiveness could be dependent not only on the coding sequence but also on surface presentation and glycosylation capacity, including availability of 9-*O*-Ac-Sia in the tissues. Together, this study positions BRO as a plausible host factor influencing OC43 host range and provides insights into understanding historical and future spillover potential within the *embecovirus* lineage.

## Extended data

**Extended Data Figure 1.**
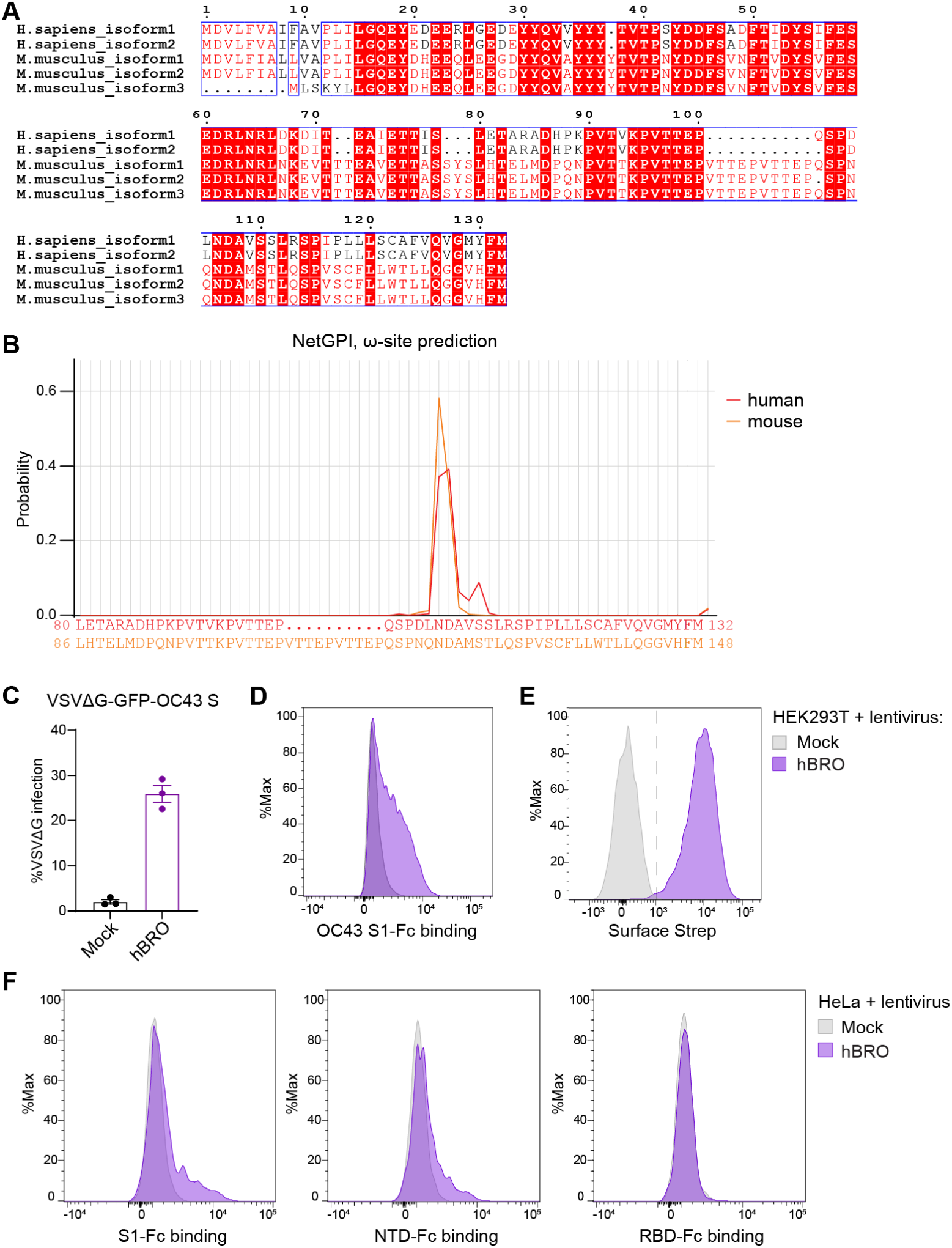
BRO is conserved in humans and mice. (A) Multiple alignment of human and mouse BRO isoforms. (B) GPI anchoring motif on human and mouse BRO was predicted using NetGPI (version 1.1). (C-E) HEK293T cells stably expressing Strep-tagged BRO were generated. (C) Cells were infected with VSVG pseudotyped with OC43 spike and GFP signal was measured at 24 hpi by flow cytometry by counting the fraction of GFP^+^ cells (n=3). (D) Flow cytometry analysis of binding of OC43 S_1_-Fc to BRO-expressing cells. (E) Surface expression of Strep tag was measured by flow cytometry. (F) OC43 spike S_1_ subunit, NTD, or RBD-Fc binding to HeLa cells stably expressing BRO was measured by flow cytometry.

**Extended Data Figure 2.**
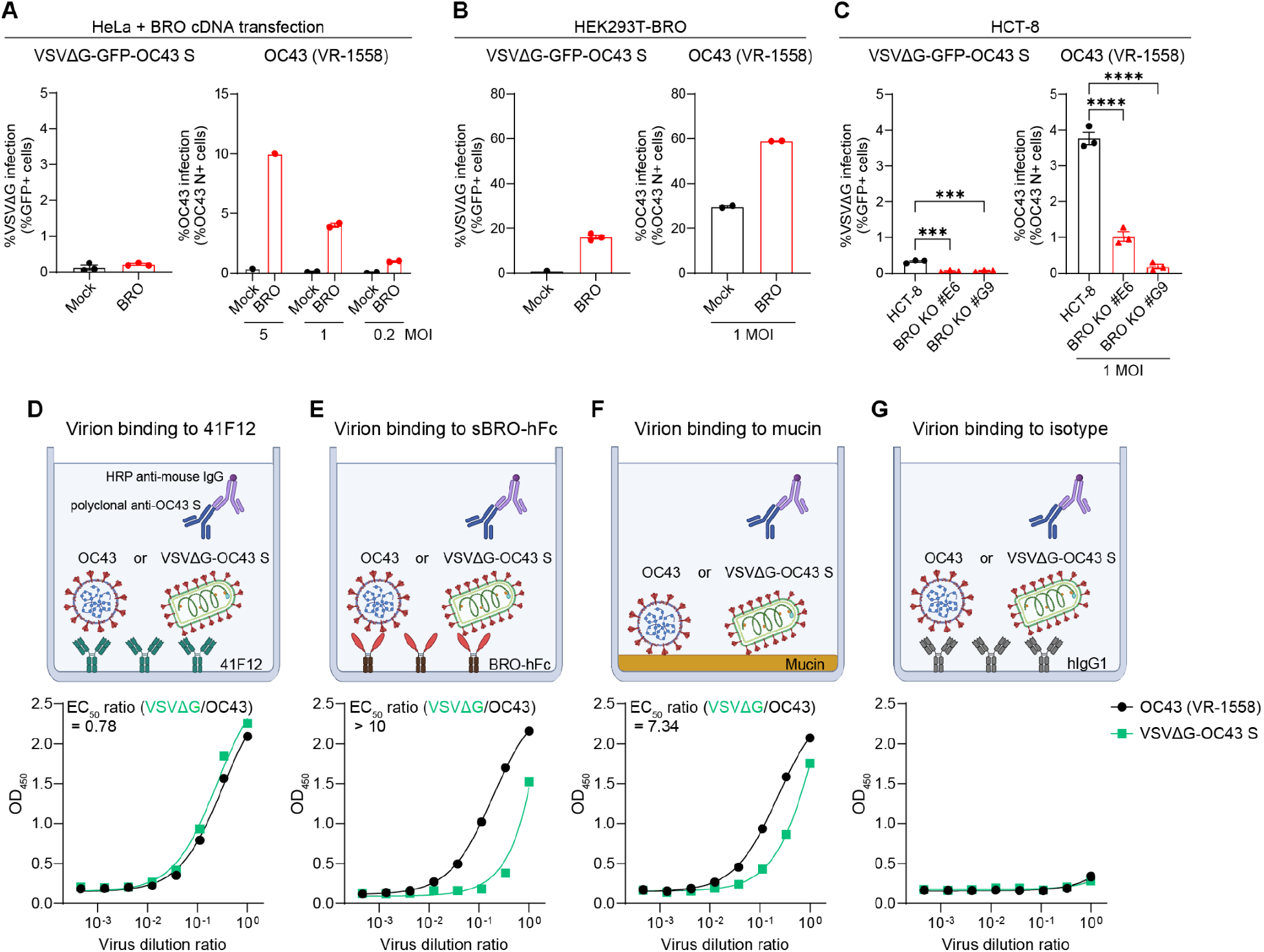
OC43 S-pseudotyped virus shows reduced BRO usage. (A-C) HeLa cells transfected with BRO cDNA (A), HEK293T-BRO stable cells (B), or WT and BRO clonal KO HCT-8 cells (C) were infected with VSVΔG pseudotyped with OC43 S or OC43 (VR-1558) at indicated MOI. GFP expression in VSVΔG-infected cells or OC43 N expression in OC43-infected cells were measured by flow cytometry. (D-G) Binding of OC43 (VR-1558) or VSVΔG-OC43 S virion to monoclonal anti-OC43 S clone 41F12 (D), soluble BRO-hFc fusion protein (E), mucin (F), or isotype control hIgG1 (G) was measured by ELISA. Serially diluted virions were added to the plate coated with indicated protein or mucin, and detected with polyclonal anti-OC43 S serum and HRP-conjugated secondary antibody. Ratio of half-maximal effective concentrations (EC50; VSVΔG/OC43) are shown.

**Extended Data Figure 3.**
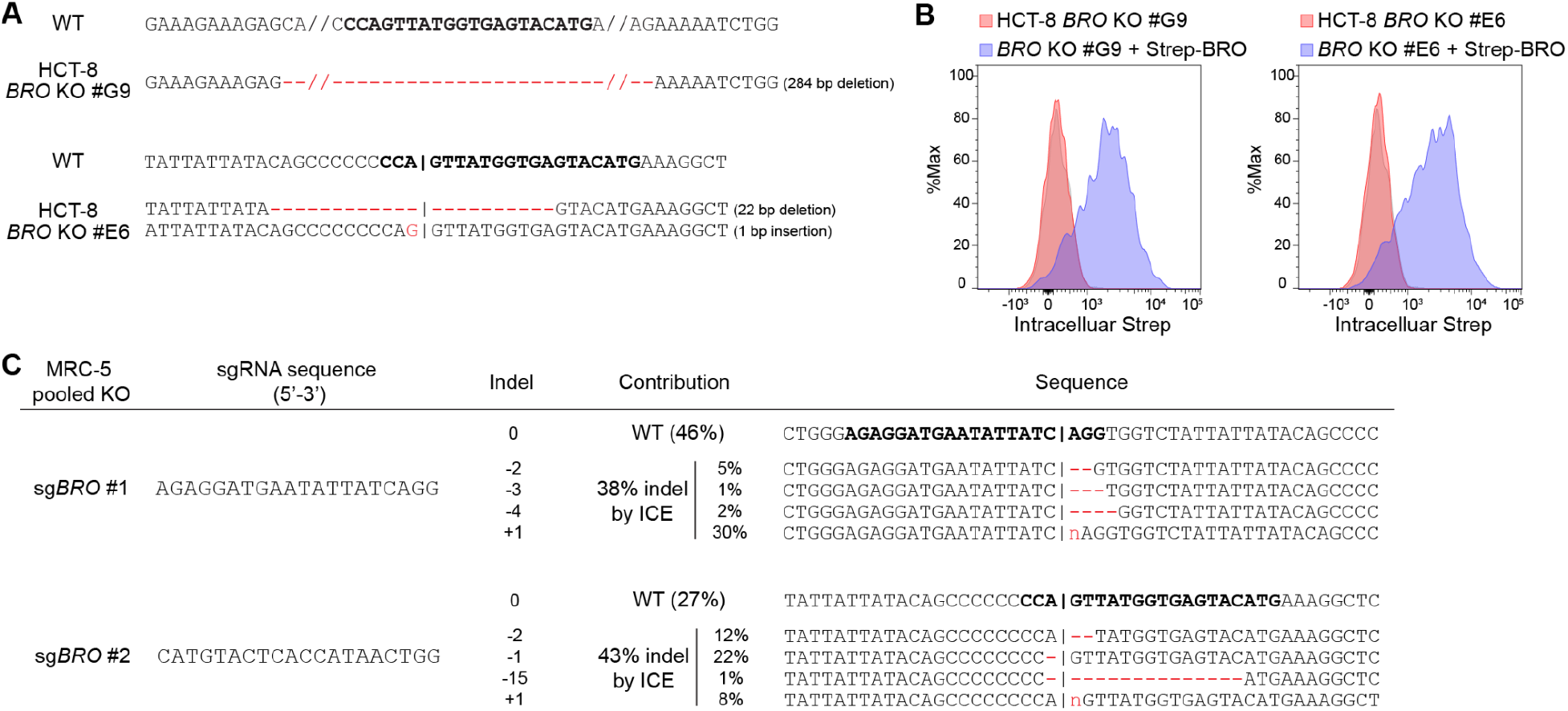
Characterization of *BRO* KO cells. (A) Edited genomic DNA isolated from HCT-8 clonal KO cells was analyzed by TOPO cloning and Sanger sequencing. (B) Intracellular Strep expression in HCT-8 *BRO* clonal KO cell lines and KO cells complemented with Strep-BRO was measured by flow cytometry. (C) Edited genomic DNA isolated from MRC-5 pooled KO cells was analyzed by Sanger sequencing and ICE analysis. Percent insertion/deletion (% indel) and contributions of each indel are shown. sgRNA sequence is highlighted in bold.

**Extended Data Figure 4.**
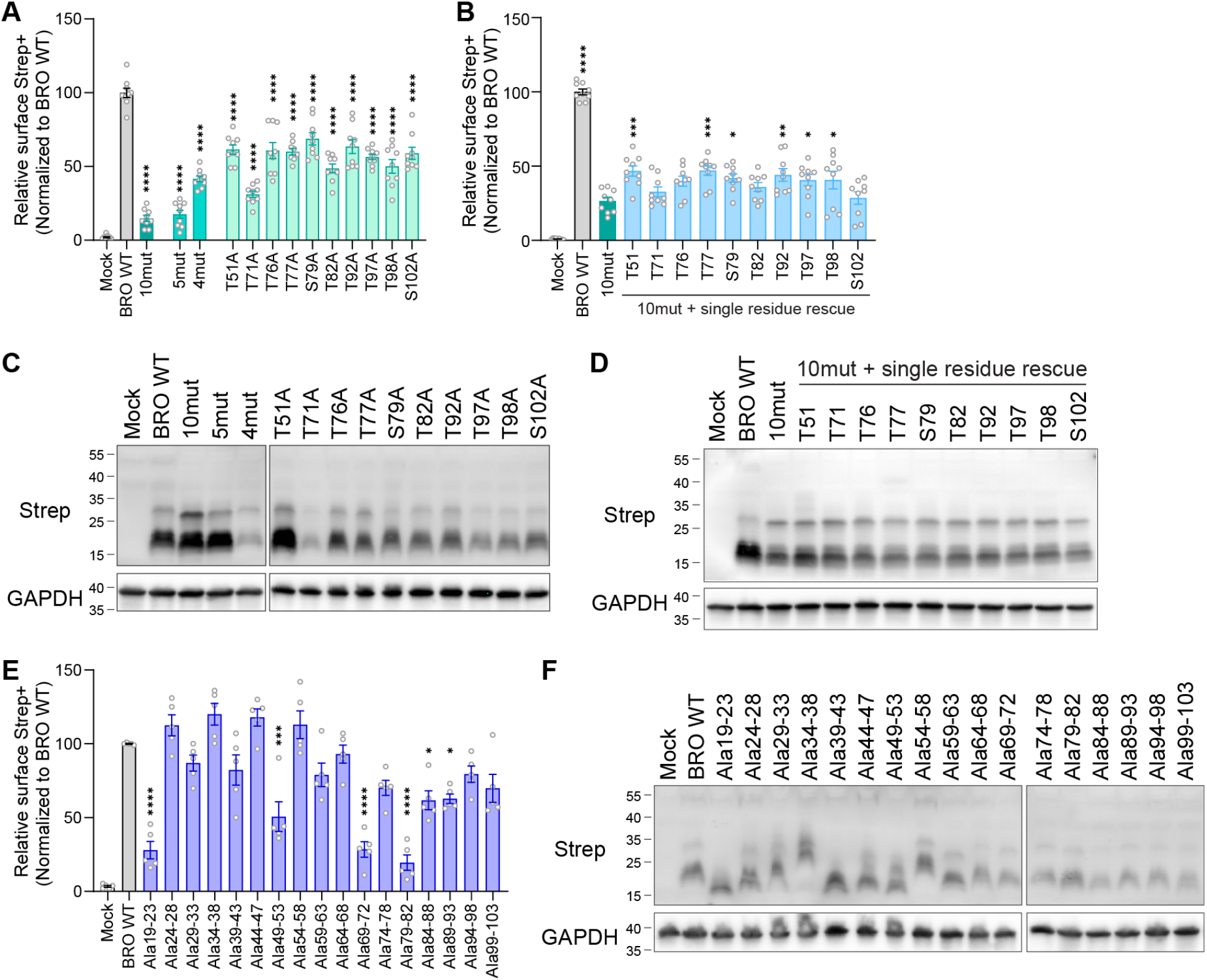
Surface and intracellular expression of BRO mutants. (A, B) HeLa cells were transfected with indicated 2xStrep-BRO mutants and surface Strep tag expression was measured by flow cytometry (n=9 from three independent experiments). (A) Residues predicted as *O*-glycosylation sites were substituted with alanine either individually or in combination. (B) Single residues of 10mut were rescued back from alanine to threonine or serine as indicated. (C, D) HeLa cells were transfected with indicated 2xStrep-BRO glycosylation mutants (C) or single glycosylation site rescue mutants (D) and Strep-BRO expression was measured by western blotting. (E, F) HeLa cells were transfected with indicated 2xStrep-BRO alanine scanning mutants. Numbers indicate amino acid residues substituted by alanine. (E) Surface Strep tag expression was measured by flow cytometry (n=5 from three independent experiments). (F) Strep-BRO expression was measured by western blotting. (A, B, E) Data represent mean ± SEM. Data were analyzed by one-way ANOVA with Tukey’s multiple comparisons test and p-values compared to BRO WT samples are shown. \**p* < 0.05; \*\**p* < 0.01; \*\*\**p* < 0.001; \*\*\*\**p* < 0.0001.

**Extended Data Figure 5.**
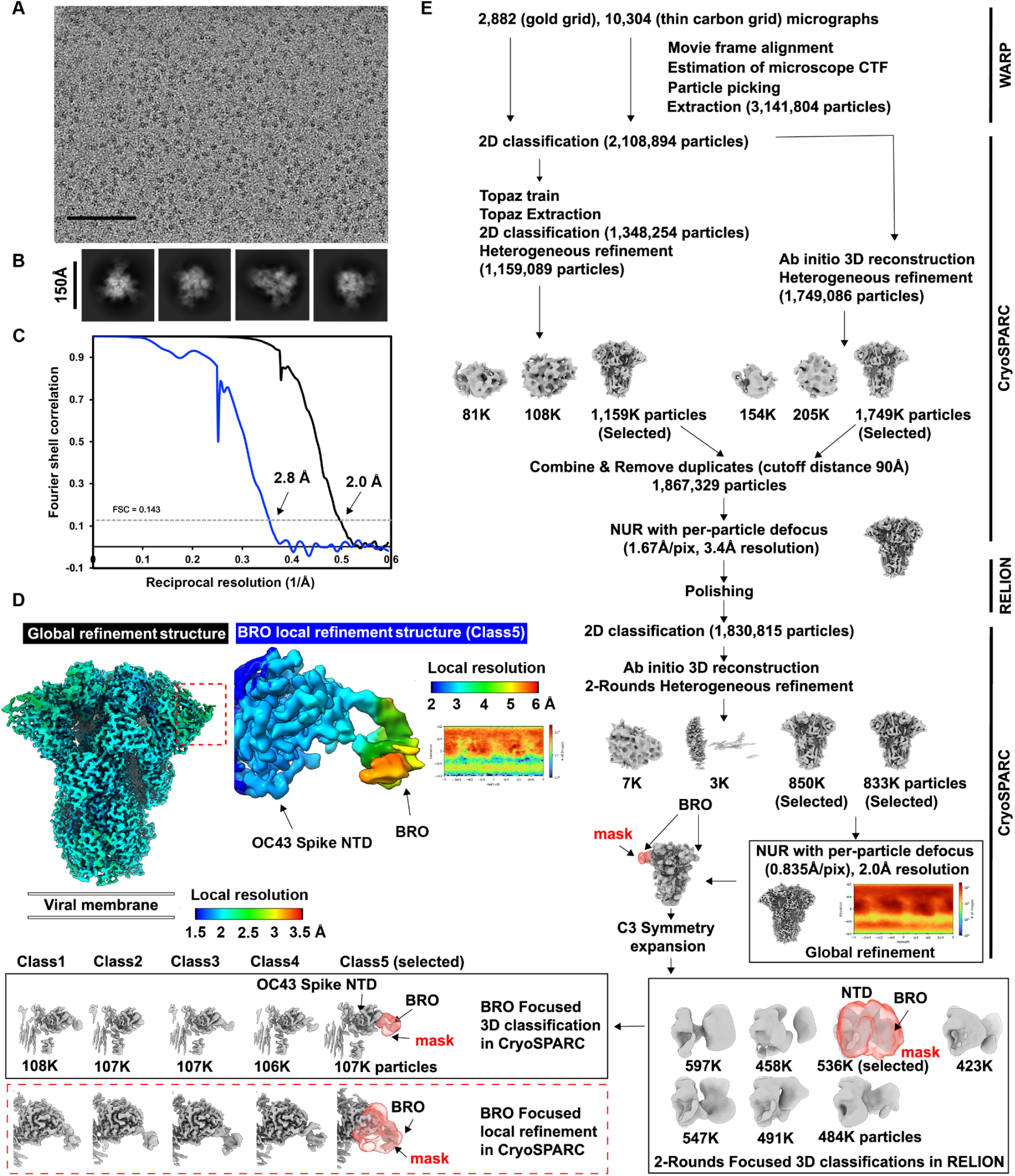
Data processing of the BRO-bound OC43 S cryoEM data set. (A-B) Representative electron micrograph and 2D class averages of BRO-bound OC43 S embedded in vitreous ice. Scale bars: 100 nm (A) and 150 Å (B). (C) Gold-standard Fourier shell correlation curves for the global refinement reconstruction (black) and BRO focused local refinement reconstruction (blue). The 0.143 cutoff is indicated by a horizontal dashed line. (D) Local resolution estimation of two structures calculated using CryoSPARC and plotted on the sharpened map of global and unsharpened map of local refinement. The red box indicated the location of the BRO focused local refinement map. (E) Data processing flowchart. CTF: contrast transfer function; NUR: non-uniform refinement. The angular distribution of particle images calculated using CryoSPARC is shown as a heat map.

**Extended Data Figure 6.**
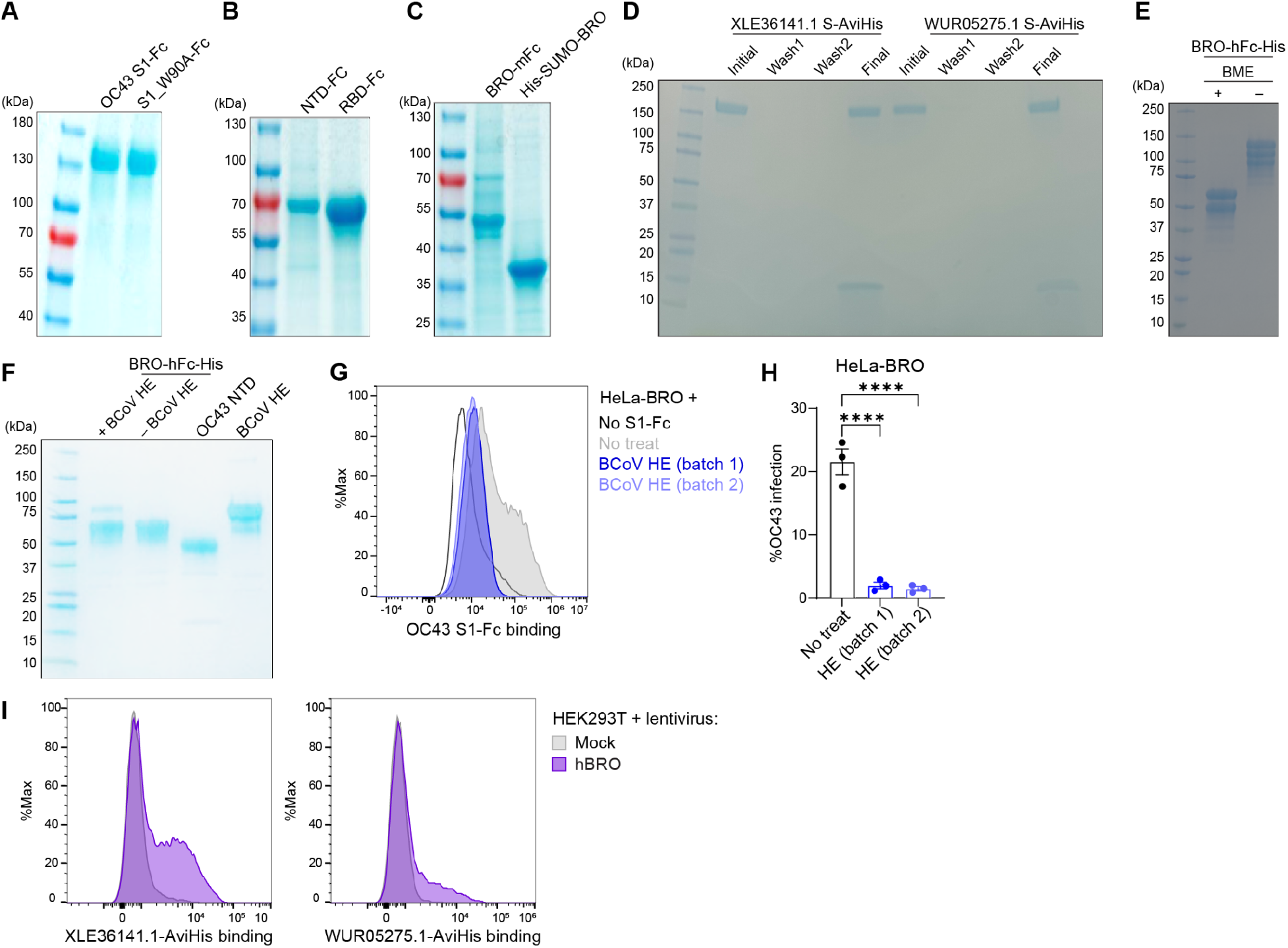
Purification of recombinant proteins used in this study. (A-F) Purified recombinant proteins analyzed by SDS-PAGE. Gels were stained with coomassie brilliant blue. (A) Purified recombinant OC43 S_1_ protein WT or sialic acid-binding residue (W90) mutated to alanine, fused with human IgG1 Fc. (B) Purified OC43 NTD and RBD protein fused with human IgG1 Fc. (C) Purified human BRO ectodomain protein fused with mouse IgG2B Fc or His-SUMO tag. (D) Biotinylated S protein of OC43 clinical isolates XLE36141.1 or WUR05275.1 with AviHis tag. (E) Purified human BRO ectodomain protein fused with human IgG1 Fc and His tag, with or without beta-mercaptoethanol (BME) treatment. (F) Recombinant proteins used for BLI analysis. BRO-hFc protein with or without BCoV HE treatment, OC43 NTD, and BCoV HE protein. (G) Flow cytometry analysis of binding of OC43 S_1_-Fc to HeLa cells stably expressing BRO pretreated with BCoV HE. (H) HeLa cells stably expressing BRO were pretreated with BCoV HE and infected with OC43 at 0.2 MOl. Intracellular OC43 N expression was measured by flow cytometry (n=3). Data represent means ± SEM and were analyzed by one-way ANOVA with Dunnett’s multiple comparisons test. ****p < 0.0001. (I) Flow cytometry analysis of binding of OC43 clinical isolates S to HEK293T cells stably expressing BRO.

**Extended Data Figure 7.**
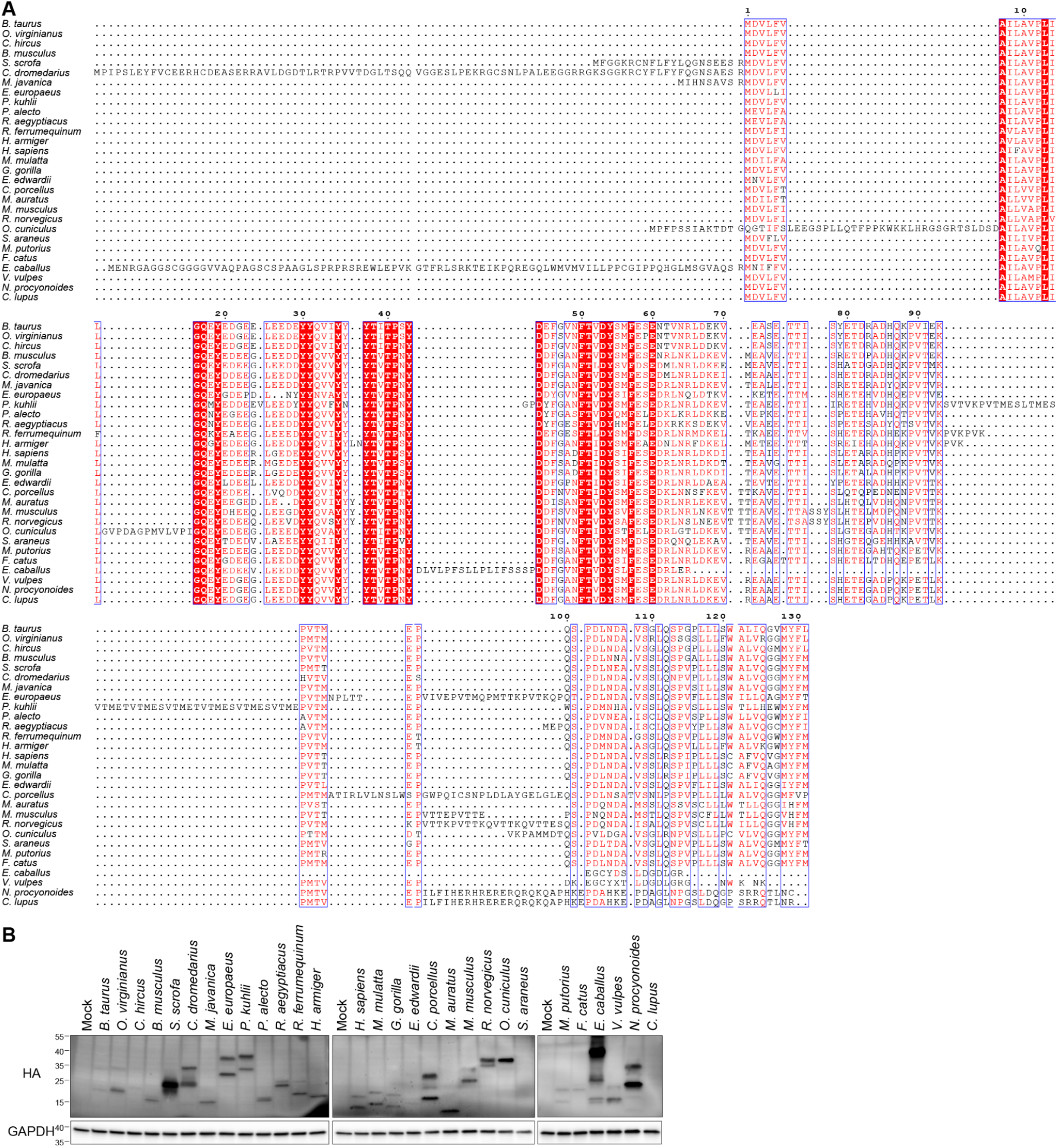
Western blot analysis and sequence alignment of mammalian BRO orthologs. (A) Amino acid sequence alignment of mammalian BRO orthologs. Sequences were aligned using MUSCLE and visualized using ESPript 3.0. (B) HEK293T cells were transfected with indicated HA-tagged BRO orthologs and HA expression was measured by western blotting.

**Extended Data Table 1.**
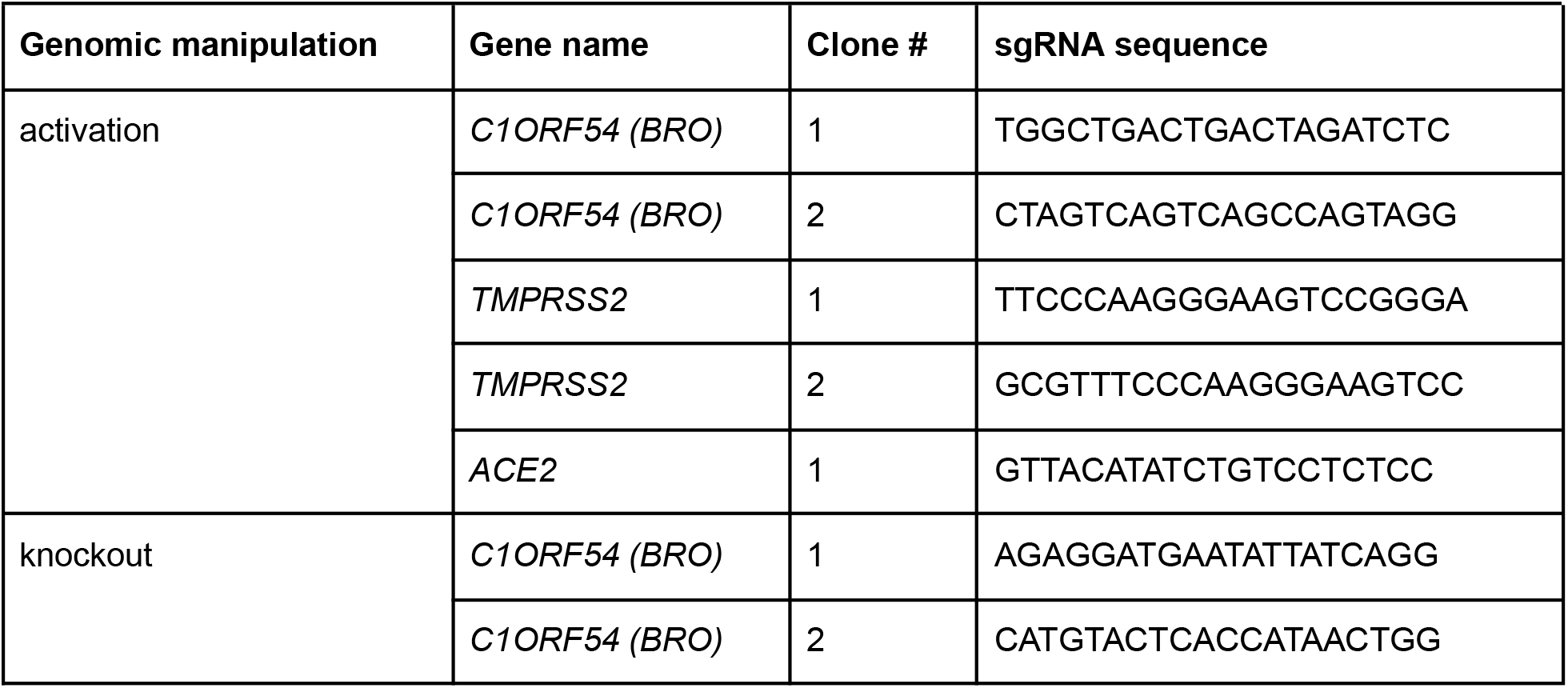
List of sgRNAs used in this study.

**Extended Data Table 2.**
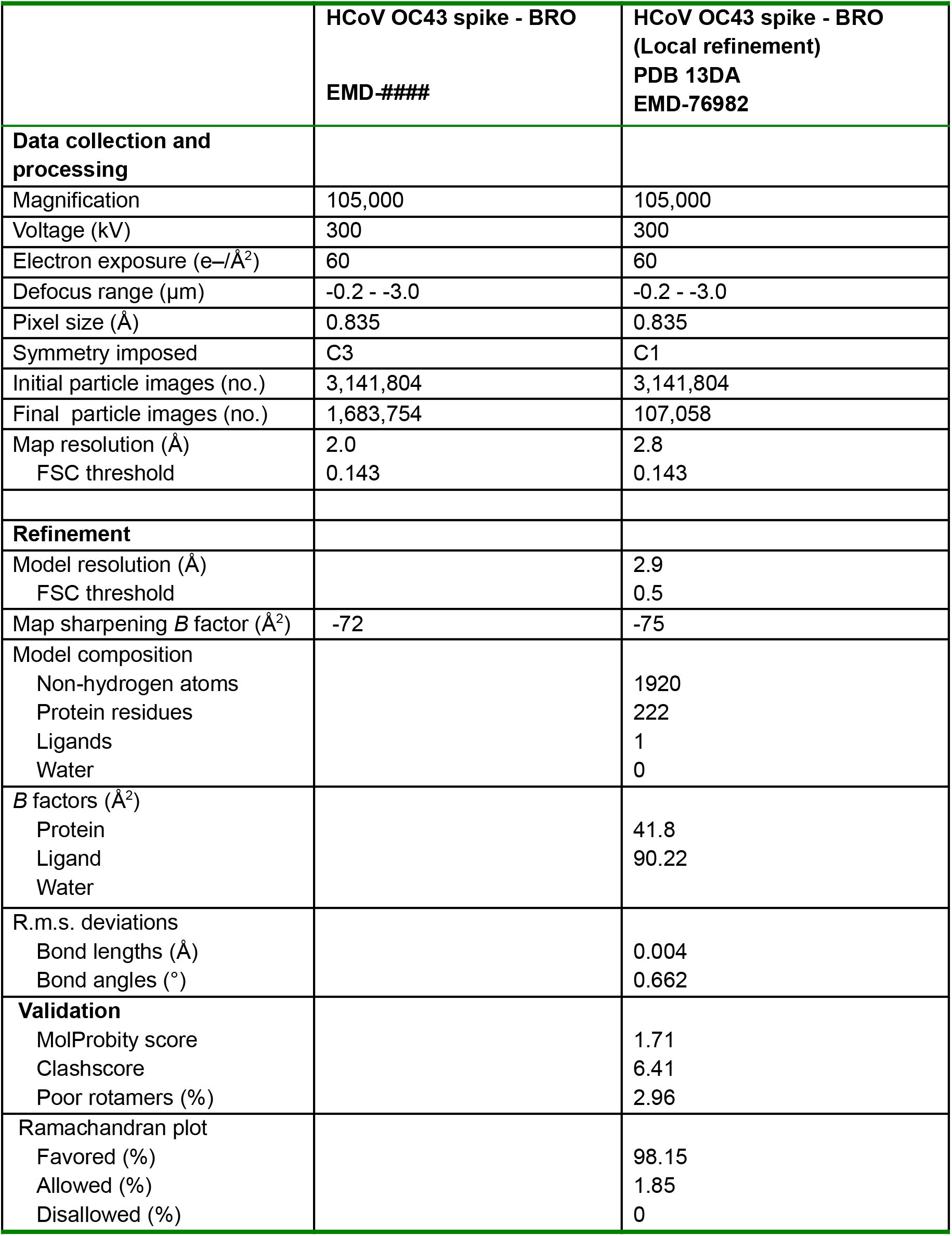
Cryo-EM data collection, refinement and validation statistics.

|  | **HCoV OC43 spike - BRO<br>EMD-####** | **HCoV OC43 spike - BRO<br>(Local refinement)<br>PDB 13DA<br>EMD-76982** |
|---|---|---|
| **Data collection and processing** |  |  |
| Magnification | 105,000 | 105,000 |
| Voltage (kV) | 300 | 300 |
| Electron exposure (e–/Å<sup>2</sup>) | 60 | 60 |
| Defocus range (µm) | -0.2 - -3.0 | -0.2 - -3.0 |
| Pixel size (Å) | 0.835 | 0.835 |
| Symmetry imposed | C3 | C1 |
| Initial particle images (no.) | 3,141,804 | 3,141,804 |
| Final particle images (no.) | 1,683,754 | 107,058 |
| Map resolution (Å) | 2.0 | 2.8 |
| FSC threshold | 0.143 | 0.143 |
| **Refinement** |  |  |
| Model resolution (Å) |  | 2.9 |
| FSC threshold |  | 0.5 |
| Map sharpening *B* factor (Å<sup>2</sup>) | -72 | -75 |
| Model composition |  |  |
| Non-hydrogen atoms |  | 1920 |
| Protein residues |  | 222 |
| Ligands |  | 1 |
| Water |  | 0 |
| *B* factors (Å<sup>2</sup>) |  |  |
| Protein |  | 41.8 |
| Ligand |  | 90.22 |
| Water |  |  |
| R.m.s. deviations |  |  |
| Bond lengths (Å) |  | 0.004 |
| Bond angles (°) |  | 0.662 |
| **Validation** |  |  |
| MolProbity score |  | 1.71 |
| Clashscore |  | 6.41 |
| Poor rotamers (%) |  | 2.96 |
| Ramachandran plot |  |  |
| Favored (%) |  | 98.15 |
| Allowed (%) |  | 1.85 |
| Disallowed (%) |  | 0 |

## Methods

### Cell culture

HEK293T (CRL-3216), HeLa (CCL-2), A375 (CRL-1619), MRC-5 (CCL-171), and HCT-8 (HRT-18; CCL-244) cells were purchased from ATCC. 3T12 cells are a gift of Herbert “Skip” Virgin. Cells were cultured in Dulbecco’s Modified Eagle Medium (DMEM, Gibco, 11995-081) supplemented with 10% fetal bovine serum (FBS; Biowest, S1620) and 2.5% HEPES (Gibco, 15630-080) at 37°C, 5% CO_2_ and detached using 0.05% trypsin-EDTA with phenol red (Gibco, 25300-120). For A375, HCT-8, and MRC-5 cells, 5 µg/mL blasticidin (Gibco, A1113903) and 1∼2 µg/mL puromycin (Gibco, A1113803) were added as appropriate. For HeLa and 3T12 cells, 100 µg/mL zeocin (Thermo Fisher Scientific, R25001) were added as appropriate. Expi293F and Expi-CHO cells (Thermo Fisher Scientific) were grown in their corresponding media at 37°C, with agitation at 110 rpm, in an 8% CO2 atmosphere and in plastic flasks with vented caps in a CO_2_ incubator.

### Construct design

All the genes encoding proteins used in BLI and cryo-EM experiments were synthesized by GenScript, codon optimized for mammalian expression and in frame with the Kozak’s sequence to direct translation except for the plasmid encoding OC43 S ectodomain previously described^1^. Gene encoding OC43 S full-length for pseudovirus production (amino acid residues 15-1329 with a deletion of the last 24 residues, Uniprot Q696P8) was cloned into pcDNA3.1(+) using a CD5 signal peptide and at the C-terminus fused to an HA tag (YPYDVPDYA) through a GGSGGS linker.

Gene encoding HE from BCoV ectodomain (residues 1-388, AZS64192.1) was cloned into pcDNA3.1(+) was followed by a linker GGS, a thrombin cleavage site, a linker GSGSGSG and a IgG1 hFc (human Fc) tag. For the pseudoviruses entry assay, the membrane-anchored HKU1 S isolate N1 (residues 1 to 1337, YP_173238, genotype A) carrying a deletion of the last 19 residues was cloned into pcDNA3.1(+) followed by the flexible linker GGSGGS and a C-terminal HA tag.

For BLI experiments, the hBRO (human BRO) -hFc construct was synthesized and cloned into a pcDNA3.1 vector. It comprises an N-terminal signal peptide (MGILPSPGMPALLSLVSLLSVLLMGCVAETGT), BRO (residues 17-106, NP_001287969.1), a C-terminal thrombin protease cleavage site, a GGGG linker, a C-terminal hFc domain, GSGGSG linker and an octa-histidine tag. OC43-NTD-WUR05275.1 (residues 1-315) and OC43-NTD-XLE36141.1 (residues 1-314) constructs were cloned into a pcDNA3.4 vector, followed by a C-terminal GS linker, an Avi tag, a GS linker and an octa-histidine tag. For prefusion S trimer proteins used in flow cytometry, the constructs OC43-S-WUR05275.1 (residues 1-1302) and OC43-S-XLE36141.1 (residues 1-1306) ectodomains were each cloned into pcDNA3.4 followed by a foldon trimerization domain, an avi-tag and a octa-histidin tag. Both constructs contain a mutation in the furin cleavage site RRSRR_759_GGSGG and the 2P mutation (A1075P and L1076P). For hBRO-mFc protein, ectodomain of hBRO (NCBI sequence ID: NP_001287969, amino acids 17-106) was cloned in the same way with mouse IgG2b Fc (AAB59658.1, amino acids 98-335) in place of the human Fc region. For recombinant BRO protein purification in *E. coli*, a codon-optimized ectodomain of human BRO was commercially synthesized and cloned into pET-28c by Genscript, downstream of a 6xHis tag and a SUMO tag including a Gly-Gly sequence to enable cleavage by Ulp1 protease.

Sequences of human BRO isoform 1 and 2 were retrieved from GenBank (accession number NP_001287968 and NP_001287969, respectively), codon-optimized, and commercially synthesized and cloned into expression vectors pTwist-CMV or pcDNA3.1 by Twist Biosciences or Genscript with a twin Strep tag after the signal peptide and a Flag tag at the C-terminus. To generate GPI anchoring mutants, coding region for amino acids N105-L119 were deleted (delGPI) from human BRO isoform 2, or N105-M131 were replaced with a GPI anchoring motif from mouse Thy-1 protein (amino acids: GGISLLVQNTSWMLLLLLSLSLLQALDFISL; exoGPI). Potential *O*-glycosylation sites were predicted using NetOGly 4.0^47^ and substituted with alanine either individually or in combination. Alanine scanning mutants were generated by substituting five consecutive residues in BRO ectodomain with alanine. All mutants were commercially synthesized and cloned into an expression vector pTwist-CMV by Twist Biosciences.

Sequences of mammalian BRO orthologs were retrieved from GenBank (accession number shown in **Supplementary Table 2**), human codon-optimized, and commercially synthesized and subcloned into an expression vector (pUC57) with a C-terminal HA-tag by Genscript. Amino acid sequences of BRO orthologs were aligned using the MUSCLE^56^ algorithm by Mega-X^57^ (version 12.0.14) and a phylogenetic tree was generated using the maximum likelihood method. Sequence alignment was visualized using ESPript 3^58^. Presence of signal peptide and GPI anchoring motif was predicted using SignalP 6.0^59^ and NetGPI 1.1^33^, respectively.

To generate OC43 S_1_-Fc fusion proteins, a mouse Ig secretory peptide was cloned into C-terminally to the wild-type S_1_ domain for OC43 (residues 14-752, YP_009555241 (ATCC VR-759 variant), or sialic acid binding deficient mutant W90A OC43 S_1_ followed by a thrombin cleavage site, a poly-glycine linker, and human IgG1 Fc (residues 99-330, AXN93647.1) by Genscript. To generate OC43 NTD-Fc and RBD-Fc proteins, coding sequences of NTD (amino acids 16-302) and RBD (amino acids 315-612) were synthesized by GenScript and cloned into pcDNA3.1 vector, downstream of an IL-2 signal sequence and upstream of a poly-glycine linker and human IgG1 Fc.

A commercial expression plasmid was used to express TMPRSS2 with a C-terminal Flag tag (Genscript, OHU13675D).

### Recombinant protein production

To produce OC43 NTD-Fc and RBD-Fc proteins, Expi293F cells were transfected with the plasmids using the ExpiFectamine293 Transfection Kit (Thermo Fisher Scientific, A14524) and treated with Enhancer 1 and 2 after 18-22 hours, following the Expi293 user guide. Five days post-tranfection, supernatants were harvested by centrifugation and filtered using a 0.45 µm membrane to remove cell debris and precipitation. Protein G Sepharose resin (Thermo Fisher Scientific, 101243) was added to the purified supernatant, and the mixture was rocked at 4°C overnight. The solution was loaded onto Econo-Pac Chromatography Columns, which were then washed three times with PBS. The protein was eluted with 0.2 M Glycine (pH 2.0) and neutralized with an equal volume of 2 M Tris base (pH 9.0), followed by buffer exchange into PBS. Purified proteins were visualized by SDS-PAGE and Coomassie blue staining.

OC43 S_1_-Fc and W90A mutant S_1_-Fc protein was expressed in CHO cells, and purified using MabSelect SuRe LX resin by Genscript.

For BRO protein expression in *E. coli* BL21(DE3), transformed bacteria with the corresponding plasmid were induced for protein expression at an OD 600 ≈ 0.6 with isopropyl β-D-1-thiogalactopyranoside followed by sequential incubation at 16°C for 16 hours and at 25°C for 6 hours. Cells were pelleted by centrifugation at 4,000 x g and resuspended in lysis buffer composed of Tris-buffered saline (TBS; pH 7.4), 10 mM DTT, 0.5% Triton X-100, supplemented with 1 μg/mL lysozyme, 1 μg/mL DNase I, and cOmplete Protease Inhibitor (Roche). Cells were sonicated using a Digital Sonifier (Branson) at 3 x 30 second bursts, centrifuged at 20,000 x g for 20 minutes, and loaded onto a Nickel NTA column. Proteins were eluted in TBS with 500 mM imidazole, followed by buffer exchange into TBS. Purified His-SUMO-BRO protein was incubated with Ulp1 protease at 6:1 ratio at 4°C, overnight and loaded onto a Nickel NTA column to remove the cleaved His-SUMO tag. Purified BRO protein was analyzed by SDS-PAGE and Coomassie blue staining and used to immunize mice as described in a separate section.

For BLI and CryoEM experiments, hBRO-hFc and hBRO-mFc protein were produced and purified using Expi293F or ExpiCHO-S cells. Expi293F cells were grown to a density of 3 x 10^6^ cells/mL and transfected using the ExpiFectamine 293 Transfection Kit (ThermoFisher Scientific) and expression carried out for 5 days post-transfection at 37°C with 8% CO_2_. ExpiCHO-S cells were grown to a density of 6 x 10^6^ cells/mL and transfected using the ExpiFectamine CHO Transfection Kit (ThermoFisher Scientific) and expression carried out for 5 days post-transfection at 37°C with 8% CO_2_.

hBRO-mFc and hBRO-hFc were purified from clarified supernatants using HisTrap HP affinity columns (Cytiva) and washed with ten column volumes of 10 mM imidazole, 25 mM Tris pH 8.0, and 150 mM NaCl before elution with two column volumes of 500 mM imidazole, 25 mM Tris pH 8.0, and 150 mM NaCl. BCoV-HE was purified using a Protein A resin, eluted using 0.1 M citric acid pH 3.0 into 1 M Tris pH 9.0 to a final pH of 8.0 then ultimately stored in the same 25 mM Tris, 150 mM NaCl pH 8.0. Purified hBRO-mFc, hBRO-hFc and BCoV-HE were then further purified by size exclusion chromatography using a Superdex 200 Increase 10/300 GL column (Cytiva) and concentrated using centrifugal filters (Amicon Ultra) before being flash frozen.

OC43-S-YP_009555241, OC43-S-WUR05275.1 and OC43-S-XLE36141.1 ectodomain proteins were produced using Expi293F. Expi293F cells were grown to a density of 3 x 10^6^ cells/mL and transfected using the ExpiFectamine 293 Transfection Kit (ThermoFisher Scientific) and expression carried out for 5 days post-transfection at 37°C with 8% CO_2_. OC43-S ectodomain proteins were purified from clarified supernatants using HisTrap HP affinity columns (Cytiva) and washed with 150 ml volume of 10 mM imidazole, 25 mM Tris pH 8.0, and 150 mM NaCl before elution with 10ml volume of 500 mM imidazole, 25 mM Tris pH 8.0, and 150 mM NaCl. After buffer exchanging to TBS pH 8.0, OC43-S proteins were biotinylated for overnight at 4°C. The biotinylated OC43-S ectodomain proteins were then further purified by size exclusion chromatography using a Superdex 6 Increase 10/300 GL column (Cytiva) at 4°C and concentrated using centrifugal filters (Amicon Ultra) before being flash frozen.

The purified recombinant proteins were assessed by SDS-PAGE followed by Coomassie blue staining (**Extended Data Fig 6A-F**).

### Transient transfections

For transient transfection, HEK293T or HeLa cells were transfected with expression plasmids using TransIT-LT1 (Mirus Bio, MIR 2300) and Opti-MEM (Gibco, 31985-070). At 24 hours post transfection, cells were detached for downstream flow cytometry or western blotting analysis, or plated into 96-well plates for infection assays.

### Surfaceome-focused CRISPR activation screen

CRISPR activation screen for surface proteins was performed as previously described^30^. Briefly, A375-dCas cells expressing CRISPR activation sgRNAs targeting 6,011 genes were incubated with 50 µg/mL OC43 S_1_-Fc fusion protein, SARS-CoV-2 spike trimer-His protein (BEI, NR-53937), or human IgG1 isotype control (BioXCell, BE0297) for 30 min at 4°C. Cells were stained with PE-conjugated anti-human IgG antibody (Southern Biotech, 9040-09) or anti-His (Abcam, ab18184) and PE anti-mouse IgG antibody (Biolegend, 405307) as appropriate. After fixing with 4% formaldehyde, cells were sorted using FACSAria III (BD Biosciences) and the top ∼3% (fluorescence intensity) of the PE-positive cells were isolated. Genomic DNA was extracted from isolated cells and unsorted cells with QIAamp DNA Mini or Maxi kit (Qiagen, 51104, 51192).

Genomic DNA samples were PCR amplified using Titanium Taq DNA polymerase (Takara, 639209), P5 stagger primer mix (0.5 µM) and uniquely barcoded P7 primer (0.5 µM), followed by purification with AMPure XP beads (Beckman Coulter, MSPP-A63880). After sequencing on a NextSeq550 sequencer (Illumina), reads were demultiplexed and mapped to each sgRNA using Library Aligner (https://gitlab.com/buchserlab/library-aligner). The log-fold change of sgRNAs in each sample was calculated compared to the unsorted sample and the p-values were determined based on the hypergeometric distribution.

The analyzed screening results are provided as **Supplementary Table 1** and the raw FASTQ files are deposited in the NCBI Sequence Read Archive Bioproject PRJNA1422175.

### Lentivirus constructs and stable cell lines generation

For lentivirus production, HEK293T cells were transfected with a desired lentiviral plasmid along with packaging plasmids, pMD2.G (Addgene, 12259) and psPAX2 (Addgene, 12260), using TransIT-LT1 (Mirus Bio, MIR 2300). The culture media was replaced at 6 hours after transfection and the supernatant was collected, centrifuged, and filtered using a 0.45 µm filter (Millipore) at 36 hours after transfection. For transduction, cells were transduced with 1 mL of lentiviral stock, spun at 1200 x g for 90 minutes at 30°C with 1 µg/mL polybrene. The following day, the culture media was replaced and cells were selected with appropriate drugs for 7 days.

For CRISPR activation, A375 cells were transduced with pLenti-dCas9-VP64-Blast (Addgene, 61425). Individual sgRNAs targeting indicated genes were cloned into linearized pXPR_502 (Addgene, 96923) and used to transduce A375-dCas9 cells. Sequences for individual sgRNAs are provided in **Extended Data Table 1**.

For CRISPR knockout, HCT-8 and MRC-5 cells were transduced with pLX_311-Cas9 (Addgene, 96924), then transduced with pLentiGuide-Puro (Addgene, 52963) harboring individual sgRNAs. Sequences for individual sgRNAs are provided in **Extended Data Table 1**. To make BRO clonal knockout HCT-8 cells, transduced cells were first selected with 5 µg/mL blasticidin and 2 µg/mL puromycin for 7 days, then diluted and plated into 96-well plates. Clonal cells were expanded from a single cell and analyzed for the targeted genome editing. Genomic DNA was extracted from BRO knockout HCT-8 and MRC-5 cells using QIAamp Blood Mini Kit (Qiagen, 51104) and the targeted region was amplified by PCR using forward primer 5’-AAAGGGCACGGAGTAACAGG-3’ and reverse primer 5’-TCGATCTCCTGACCTCGTGA-3’. PCR products were used for TOPO cloning to confirm the edited sequences or analyzed by Sanger sequencing and ICE analysis (EditCo) for HCT-8 clones and MRC-5 cells respectively.

For stable expression of BRO, a lentiviral vector pCDH-MSCV-T2A-Puro (System Biosciences, CD522A-1) was modified to contain an EF1alpha promoter, beta-globin intron, internal ribosome entry site, and a zeocin selection marker (pCDH-EF1a-intron-IRES-Zeo). The desired 2xStrep-BRO-Flag sequences were PCR-amplified from pcDNA3.1 expression vectors and subcloned into the lentiviral vector using NEBuilder HiFi DNA assembly master mix (NEB, E2621S). Lentiviral particles production in HEK293T cells and transduction into HeLa or 3T12 cells were performed as described above.

### Generating mouse polyclonal antibodies against human BRO, and OC43 virus

Wildtype C57BL/6J (stock #000664) mice were obtained from Jackson Laboratory and housed at the Brown University facilities under specific-pathogen-free conditions with food and water ad libitum and 12-hour day-night cycle. Mice experiments were performed following a protocol approved by Brown University Institutional Animal Care and Use Committee (protocol #23-04-0002).

Adult female mice were subcutaneously immunized with 30 μg of recombinant human BRO ectodomain at day 0 and with 15 μg of the immunogen at days 14, 26, 40, and 53. Quil-A adjuvant (Invivogen, vac-quil) was mixed with the immunogen for immunization. Specific IgG levels in the serum were titrated by ELISA against recombinant BRO ectodomain. To produce polyclonal antibodies targeting OC43 spike, OC43 virus was UV-irradiated at 300 mJ/cm^2^ before use. Adult female mice were immunized with 5 x 10^6^ PFU of inactivated OC43 virus at day 0 and with 2.5 x 10^6^ PFU at days 14, 26, 40, and 53 as described for BRO immunization. Serum samples were titrated by ELISA against OC43 virion or recombinant spike proteins.

### Flow cytometry

To measure OC43 spike binding to cells expressing BRO, BRO-induced A375-dCas cells, HEK293T transfected with BRO cDNA, or HEK293T and HeLa cells stably expressing BRO were used. Cells were washed once with HBSS containing 2% FBS and incubated with 100 µg/mL S_1_-Fc, RBD-Fc, or NTD-Fc for 30 minutes at 4°C, then stained with PE anti-human IgG (Southern Biotech, 9040-09) for 30 minutes. To measure OC43 clinical isolate S trimer binding, cells were incubated with 100 µg/mL His-tagged S proteins for 30 minutes, followed by staining with anti-His antibody (Abcam, ab18184) and PE anti-mouse IgG (Biolegend, 405307). Cells were fixed with 4% paraformaldehyde for 10 minutes and resuspended in FACS buffer (1x PBS supplemented with 2% FBS and 1 mM EDTA) for flow cytometry. When indicated, cells were pretreated with 20 µg/mL bovine CoV HE protein in PBS for 30 minutes at 37°C before subjecting to OC43 S_1_-Fc binding.

To measure the expression of Strep or Flag tag on the surface, cells were stained with Strep-Tactin XT DY-649 (IBA Lifesciences, 2-1568-050) or BV421 anti-Flag (Biolegend, 637321) in FACS buffer for 30 minutes at 4°C, washed two times in FACS buffer, and fixed with 4% paraformaldehyde before being analyzed. Surface expression of endogenous human BRO after CRISPR activation in A375-dCas cells was measured using BRO-immunized mouse serum and PE anti-mouse IgG (Biolegend, 405307). To measure intracellular expressions of the tags, cells were first fixed with Cytofix/Cytoperm buffer (BD Bioscience) for 10 minutes at room temperature, followed by washing twice with Perm/Wash buffer (BD Bioscience), and stained with the same reagents as for surface staining. Cells were analyzed using Cytek Aurora spectral analyzer (Cytek Biosciences). Data was analyzed with FlowJo software.

### Pseudovirus production and entry assay

VSVΔG-GFP and VSVΔG-Luc pseudoviral particles were produced as previously described with minor modifications^60^. HEK293T cells were plated in 10-cm tissue culture dishes and transfected with plasmids encoding OC43 or HKU1 spike proteins using Lipofectamine 3000 (Invitrogen). To express CoV spike proteins for pseudotyping, an expression plasmid for SARS-CoV-2 B.1.1.7 spike (Addgene, 170451) was used as a template, and its spike coding region was replaced with the coding sequences of OC43 spike (GenBank accession number YP_009555241, amino acids 1-1336) and HKU1 spike (ADN03339, amino acids 1-1339) by Genscript. For OC43 pseudovirus, OC43 HE-encoding plasmid (Sino Biological, VG40603-UT) was co-transfected at 1:16 ratio. At 24 hours post transfection, cells were infected with replication restricted rVSVΔG∗G-GFP virus (Kerafast, EH1019-PM) at 5 MOI for 1 h at 37°C, then the media was replaced with complete media. Anti-VSV-G (Sigma, MABF2337) was added at final concentration of 1 μg/mL to neutralize residual rVSVΔG∗G. After another 24 hours, viral supernatant was harvested, cell debris were removed by centrifuging for 10 min at 1320 x g and stored at -80°C in small aliquots.

For entry assay, 2 x 10^4^ cells were plated per well in 96-well plates. The following day, the culture media was removed and cells were incubated with 100 μL of pseudoviral particles for 20∼24 h at 37°C, 5% CO_2_. Cells were detached using trypsin, fixed with 4% paraformaldehyde, and resuspended with FACS buffer. GFP-positive cells were measured by flow cytometry using Cytek Aurora spectral analyzer (Cytek Biosciences).

### Authentic OC43 virus preparation and infection

HCoV OC43 virus (ATCC VR-1558 variant) was obtained from BEI Resources (NR-56241) and passaged once in MRC-5 cells to generate a working stock. MRC-5 cells were plated in 15-cm dishes and incubated with the virus for 2 days at 33°C, 5% CO2. Cells and supernatants were freeze-thawed and centrifuged to remove the cell debris. Virus-containing supernatants were filtered and concentrated with Vivaflow 50 (Sartorius). Virus titer was determined via plaque assay. MRC-5 cells were plated in 6-well plates and infected with serial dilutions of viral stock for 1 hour at room temperature. After removing the inoculum, cells were overlaid with 1% methylcellulose (Sigma) in MEM (Gibco) supplemented with 5% FBS, 1x Pen/Strep, and 10 mM HEPES. Cells were incubated for 5 days and fixed and stained with 0.1% crystal violet/20% ethanol.

For viral infection, 2 x 10^4^ HCT-8, MRC-5, HeLa, or 3T12 cells were plated per well in 96-well plates. Next day, the media was removed and cells were incubated with virus inoculum for 2 hours at 33°C, 5% CO_2_. After removing the inoculum, cells were incubated in fresh complete media at 33°C for 24 or 48 hours and cells or supernatants were harvested to measure infectivity or virion titer.

When indicated, cells were pretreated with 0.1 or 1 U/mL Phosphatidylinositol-Specific Phospholipase C (PI-PLC; Invitrogen, P6466), 10 μM Camostat mesylate (Sigma, SML0057), 10 μM E64d (Aloxistatin; MedChemExpress, HY-100229), or 20 μg/mL recombinant bovine CoV HE for 1 hour at 37°C, washed once with PBS, then treated with virus inoculum. After 2-hour adsorption at 33°C, the inoculum was removed and fresh media was added along with the enzyme or entry inhibitors.

To measure viral infectivity, cells were fixed with Cytofix/Cytoperm buffer (BD Bioscience) for 10 minutes at room temperature, followed by washing twice with Perm/Wash buffer (BD Bioscience), and stained with anti-OC43 Nucleocapsid (N) antibody (clone 541-8F; Millipore, MAB9012) or rabbit polyclonal anti-OC43 N antibody (SinoBio, 40643-T62) for 30 minutes at 4°C. Cells were washed twice with Perm/Wash buffer and stained with PE-conjugated anti-mouse IgG (Biolegend, 405307) or anti-rabbit IgG (Jackson ImmunoResearch Laboratories, 711-116-152) for another 30 minutes. After washing, cells were analyzed using Cytek Aurora spectral analyzer (Cytek Biosciences). Data was analyzed with FlowJo software.

### OC43 virion titration by focus forming assay

To measure OC43 virion production in 3T12-BRO and BRO knockout HCT-8 or MRC-5 cells, cells were infected with OC43 for 2 hours at 33°C, then virus inoculum was replaced with complete media. Supernatant was harvested at 1, 24, or 48 hpi as indicated. 2 x 10^4^ HCT-8 cells were plated per well in 96-well plates (Agilent, 204626-100) and infected with serial dilutions of culture supernatants for 2 hours at 33°C. After removing the inoculum, cells were overlaid with 1% methylcellulose (Sigma) in MEM (Gibco) supplemented with 5% FBS, 1x Pen/Strep, and 10 mM HEPES and incubated for 5 days. Cells were fixed with 10% formaldehyde for 1 hour at room temperature, permeabilized using 0.1% Triton X-100, and blocked with 10% goat serum. Cells were stained with mouse monoclonal anti-OC43 N antibody (clone 541-8F; Millipore, MAB9012) at 4°C overnight, followed by secondary staining with AF555 anti-mouse IgG (Thermo Fisher Scientific, A32727) for 1 hour. The number of total cells and OC43 N-positive cells were analyzed using Cytation 5 (Biotek) and the focus forming units per mL were calculated.

### Western blot

Cells were lysed in RIPA Lysis and Extraction buffer (Thermo Fisher Scientific) with cOmplete Protease Inhibitor (Roche) for 10 minutes at 4°C and spun down at 18,000 x g for 30 minutes. Lysate was mixed with the SDS sample buffer and boiled for 10 minutes. For PI-PLC treatment, cells were incubated with 10 U/mL PI-PLC (Invitrogen, P6466) for 1 hour at 37°C and washed once with PBS, followed by lysis using RIPA buffer. Protein samples were separated by SDS-PAGE using 4–20% Mini-PROTEAN TGX Stain-Free Protein Gels (Bio-Rad), transferred to 0.45 µm PVDF membrane (Bio-Rad), blocked with Intercept (TBS) Blocking Buffer (Li-Cor), and probed with indicated antibodies. Signals were visualized with the ChemiDoc MP Imaging System (Bio-Rad).

Antibodies used for western blotting include: anti-Strep II (Genscript, A01732), anti-FLAG (Rockland Immunochemicals, 200-350-383), anti-HA (Biolegend, 901533), DyLight800-conjugated anti-GAPDH (Bio-Rad, MCA4739), and StarBright Blue 700-conjugated anti-mouse IgG (Bio-Rad, 12004158).

### Neutralization assays and RT-qPCR

2 x 10^4^ HCT-8 cells were plated per well in 96-well plates one day prior to infection. To test neutralizing activity of recombinant BRO-Fc, OC43 virus (100 PFU) was pre-incubated with serial dilutions of BRO-hFc or isotype control for 1 hour at 33°C. Cells were treated with virus-protein mixture for 2 hours at 33°C for virus adsorption. For neutralization of BRO, cells were pretreated with anti-BRO mouse serum for 1 hour at 37°C, followed by treatment with virus inoculum (500 PFU) for 2 hours at 33°C for virus adsorption. In both assays, virus inoculum was removed after the adsorption period and replaced with fresh complete media. Infected cells were incubated at 33°C for 48 hours and supernatant was collected for RNA analysis.

RNA was isolated from stool using Quick-RNA Viral 96 kits (Zymo Research, R1041) according to the manufacturer’s protocol and cDNA was synthesized using ImPromII reverse transcriptase system (Promega, PAA3803). Quantitative PCR was performed with AmpliTaq Gold DNA polymerase (Applied Biosystems, 4311818) and following primers and probe targeting OC43 N (forward primer 5’-CGATGAGGCTATTCCGACTAGGT-3’, reverse primer 5’-CCTTCCTGAGCCTTCAATATAGTAACC-3’, probe /56-FAM/TCCGCCTGG/ZEN/CACGGTACTCCCT/3IABkFQ/; synthesized at IDT), using a standard curve for determination of absolute viral genome copies.

### Human bronchial epithelial cell culture, differentiation, and neutralization

Human bronchial epithelial cells (HBECs) were purchased from the American Type Culture Collection (ATCC) and initially expanded in PneumaCult-Ex Plus Medium (STEMCELL Technologies, 05040) supplemented with 1% penicillin–streptomycin (Gibco, #15140122). Cells were maintained in T75 vented, tissue culture–treated flasks and culture medium was replaced every two days until cells reached 80–90% confluency.

For differentiation at the air–liquid interface (ALI), HBECs were gently trypsinized and seeded onto the apical surface of collagen-coated Transwell inserts (STEMCELL Technologies, #38024) at a density of 5 × 10^4^ cells per insert in 100 µL of PneumaCult-Ex Plus Medium. Both apical and basal compartments were fed every two days until the cells reached confluence. Upon confluence, the apical medium was removed to establish ALI conditions, and PneumaCult-ALI Maintenance Medium (STEMCELL Technologies, 05001) was added exclusively to the basal compartment. Basal medium was changed every two days, and starting one week after exposure to air, the apical surface was washed twice weekly warm PBS to remove excess mucus. Cells were maintained under ALI conditions for up to 21 days to allow full differentiation. All cultures were maintained at 37°C in a humidified atmosphere containing 5% CO_2_.

For neutralization of BRO, cells were washed at the apical side and pretreated with anti-BRO mouse serum (made in-house) for 30 minutes at 37°C, followed by treatment with virus inoculum (10^6^ PFU) for 2 hours at 33°C for virus adsorption. After the adsorption, virus inoculum was removed and cells were washed three times with PBS. Infected cells were incubated at 33°C for 48 hours and trypsinized for RNA analysis.

### Immunofluorescent staining

Human bronchial epithelial cells (HBECs) differentiated at air–liquid interface (ALI) were used. The transwell membranes were fixed in 4% paraformaldehyde for 15 minutes at 4°C and washed three times with 1x PBS, then embedded in 30% sucrose (Sigma, S3089) solution at 4°C overnight. Later, the membranes were submerged in a mixture of 100% OCT tissue freezing medium (IMEB, DFM-2) and 30% sucrose at 1:1 ratio for 2 hours at room temperature. Later, the membrane was frozen with 100% OCT for 15 minutes on a dry ice ethanol slurry. The cryosections were prepared by using a Leica CM1860 cryostat and each cryosection was 8 µm.

To perform immunofluorescence staining, the slides were gently rinsed with UltraPure Distilled water (Invitrogen, 10977-015) and then permeabilized with 0.1% Triton X-100 (Sigma, T8787). The sections were blocked in 10% goat serum (Abcam, AB7481) in PBS for 1 hour at room temperature. The sections were incubated with anti-BRO primary antibody diluted at 1:200 with 10% goat serum in PBS overnight at 4°C, followed by incubation of goat anti-mouse secondary antibody (Invitrogen, A32728) at 1:2000 dilution for 1 hour at room temperature. Coverslips were then mounted with ProLong Glass Antifade Mountant with NucBlue (Invitrogen, P36981). The confocal images were acquired using an Olympus FV3000 confocal microscope.

### Biolayer interferometry assay (BLI)

To assess the interaction between OC43 NTDs and hBRO we performed binding experiments using BLI on an Octet Red instrument (Sartorius) operated at 30°C with 1000 rpm shaking. Biotinylated OC43 S NTDs from YP_009555241.1, XLE36141.1 and WUR05275.1 strains were immobilized at 3 µg/mL to reach 1 nm shift in 10X kinetics buffer (Pall) to streptavidin (SA) biosensors (Sartorius) pre-hydrated in 10X kinetics buffer for 10 minutes prior the experiment. To stabilize and remove unbound proteins, biosensors were subsequently dipped into 10X kinetics buffer, after which were dipped in a solution containing 1 µM purified hBRO. The dissociation phase was followed by dipping the tips into 10X kinetics buffer. To assess the effect of BCoV-HE on hBRO binding to OC43 NTD, 22 µg of hBRO was treated with 2.4 µg of BCoV-HE during 30 minutes at 37°C. Immediately after, SA biosensors loaded with biotinylated YP_009555241.1 OC43 NTD were dipped in a solution containing 1.5 µM of hBRO treated or not with BCoV-HE. Subsequently the loaded tips were dipped into 10X kinetics buffer during the dissociation phase. Data were plotted in Graphpad Prism (v.10.2.2).

### Cryo-electron microscopy data collection, processing and model building

The hBRO bound HCoV OC43 spike ectodomain complex was prepared by mixing a 5:1 molar ratio of the hBRO-mFc with HCoV OC43 spike ectodomain (YP_009555241.1, ATCC VR-759 variant) before incubation for one hour on ice. Specimen vitrification followed two methods. 3 µL of 0.46 mg/ml complex was applied onto freshly glow discharged a lacey carbon grid with a home-made continuous thin carbon layer prior to plunge freezing using a vitrobot MarkIV (ThermoFisher Scientific) with a blot force of -1 and 2.5 sec blot time at 100% humidity and 22°C. 3 µL of 0.46 mg/mL complex was added to the glow discharged side of R 2/2 UltrAuFoil grids^61^ and 1 µL was added to the back side before plunging into liquid ethane using a GP2 (Leica) with 6 sec blot time. The data were acquired using an FEI Titan Krios transmission electron microscope operated at 300 kV and equipped with a Gatan K3 direct detector and Gatan Quantum GIF energy filter, operated in zero-loss mode with a slit width of 20 eV. Automated data collection was carried out using Leginon^62^ at a nominal magnification of 105,000x with a pixel size of 0.835 Å. The dose rate was adjusted to 10.5 counts/pixel/s, and each movie was acquired in counting mode fractionated in 100 frames of 40 ms. A total 13,186 movies were collected with a defocus range between -0.2 and -3 μm and stage tilt angle of 0°, 25° and 30°. Movie frame alignment, estimation of the microscope contrast-transfer function parameters, particle picking, and extraction were carried out using Warp^63^. Particles were extracted with a box size of 260 pixels with a pixel size of 1.67 Å. The reference-free 2D classification was performed using cryoSPARC^64^ to select well-defined particle images. Initial model generation was carried out using ab-initio reconstruction in cryoSPARC^64^ and the resulting maps were used as references for heterogeneous 3D refinement. Particles belonging to classes with the best resolved BRO bound HCoV OC43 spike ectodomain complex density were selected. To further improve the data, the Topaz^65^ model was trained on Warp-picked particle sets belonging to the best classes after 2D classification and particles picked using Topaz were extracted and subjected to 2D classification and heterogenous 3D refinements. The two particle sets from the Warp and Topaz picking strategies were merged and duplicate particles were removed using a minimum distance cutoff of 90 Å. The 3D refinement was carried out using non-uniform refinement with per particle defocus in cryoSPARC^66^. The dataset was transferred from cryoSPARC to Relion using the pyem program package^67^ and particle images were subjected to the Bayesian polishing procedure implemented in Relion^68^ during which particles were re-extracted with a box size of 512 pixels and a pixel size of 0.835 Å. After polishing, the reference-free 2D classification was performed using cryoSPARC. To further improve the map quality, the four models generated by ab-initio reconstruction were used as references for heterogeneous 3D refinement in cryoSPARC. The final 3D refinements were carried out using non-uniform refinement along with per-particle defocus refinement in cryoSPARC^66^ to yield the final reconstruction at 2.0 Å resolution comprising 1,683,754 particles. To improve the density of the Spike NTD-BRO, the particles were symmetry expanded and subjected to two rounds of focus 3D classification without refining angles and shifts using a soft mask encompassing the BRO and a tau value of 10 in Relion^69^. Particles belonging to class with the best resolved BRO density were selected and then subjected to focused 3D classification in CryoSPARC, and followed by local refinement using CryoSPARC. The selected class5 dataset contained 107,058 particles was used for the final local refinement^64^ with a soft mask comprising BRO domain and OC43 NTD resulting in a 2.8 Å resolution reconstruction. Reported resolutions are based on the gold-standard Fourier shell correlation (FSC) of 0.143 criterion and Fourier shell correlation curves were corrected for the effects of soft masking by high-resolution noise substitution^70,71^. Local resolution estimation was carried out using cryoSPARC. Map sharpening in cryoSPARC, ARdecon^72^, and EMready^73^ was performed to assist model building. UCSF Chimera^74^, Coot^75^, and Phenix^76^ were used to fit, build, and refine the model using the sharpened and unsharpened cryo-EM maps. Validation used Phenix, Molprobity^77^, EMRinger^78^ and Privateer^79^.

### ELISA binding assay

ELISA assays were performed using BD OptEIA reagent set B (BD Biosciences, 550534) and 96-well EIA/RIA plates (Corning, 3361).

Live OC43 (strain VR-1558) and VSVΔG pseudotyped with OC43 spike were used for ELISA assays after UV irradiation at 300 mJ/cm^2^. To titrate the amount of spike proteins, plates were coated with serially diluted virus stocks at 4°C overnight. After 1 hour blocking, the plates were sequentially incubated with in-house polyclonal anti-OC43 spike mouse antibody and HRP-conjugated anti-mouse IgG (Southern Biotech, 1030-05) for 1 hour at room temperature. TMB substrate was added to the plates and then quenched with stop solution after 30 minutes. Absorbance at 450 nm was recorded with a Synergy HT microplate reader (Biotek).

To measure binding of virions to soluble BRO, plates were coated with 0.1 μg of BRO-hFc protein, monoclonal anti-OC43 clone 41F12^36^, human IgG1 isotype control, or 0.5 μg of bovine submaxillary mucin (Sigma, M3895) at 4°C overnight. After 1 hour blocking, the plates were incubated serially diluted live OC43 and VSVΔG-OC43 pseudovirus for 2 hours at room temperature. Then the plates were incubated with anti-OC43 spike mouse antibody, HRP-conjugated anti-mouse IgG, and TMB substrate and analyzed as described above.

To examine the glycan dependency of OC43 spike S_1_-BRO binding, plates were coated with 1 μg of BRO-mFc protein at 4°C overnight, followed by 1 hour blocking. The plates were incubated with 20 mU/mL neuraminidase (Sigma, N2876) at 37°C for 3 hours and washed three times. Then the plates were treated with serially diluted OC43 WT or W90A mutant spike S_1_-Fc proteins for 1 hour at room temperature, and incubated with HRP-conjugated anti-human IgG (Southern Biotech, 2014-05). TMB substrate was added to the plates and analyzed as described above.

### Statistical analysis

Statistical significance was determined using GraphPad Prism 10 software. Experiments were analyzed by one-way ANOVA with Kruskal-Wallis test, Dunnett’s multiple comparisons test, or Tukey’s multiple comparisons test and two-tailed unpaired Mann-Whitney test as indicated.

## Data availability

All data reported here are available from the corresponding authors upon request. CRISPR screening data generated in this study are provided as Supplementary Table 1 and the raw FASTQ files can be found at the NCBI Sequence Read Archive (SRA) under Bioproject PRJNA1422175. The cryoEM structures and maps have been deposited to the Protein Data Bank and Electron Microscopy Data Bank with accession codes PDB ID 13DA and EMD-76982.

## Acknowledgement

We acknowledge the Brown University Flow Cytometry and Sorting Facility, the Genomics facility, CARE animal facility, and the Leduc Bioimaging facility for help with critical analysis; the Washington University Genome Engineering & Stem Cell Center for CRISPR reagents. Schematic illustrations were created with BioRender.com.

Funding information: This work was supported by the National Institutes of Health grant R01 AI183155 (S.L.), R21 AI173821 (S.L., C.B.W.), P01AI167966 (D.V.), DP1AI158186 (D.V.) and 75N93022C00036 (D.V.), the Smith Family Awards Program for Excellence in Biomedical Research (S.L.), the Charles H. Hood Foundation Child Health Research Award (S.L.), an Investigators in the Pathogenesis of Infectious Disease Awards from the Burroughs Wellcome Fund (D.V.), and the University of Washington Arnold and Mabel Beckman cryoEM center. D.V. is an Investigator of the Howard Hughes Medical Institute and the Hans Neurath Endowed Chair in Biochemistry at the University of Washington.

## Author Contributions

Conceptualization: J.S., Y.-J.P., D.V., S.L.

Methodology: J.S., Y.-J.P., M.A.T., L.Z., S.M., D.V., S.L.

Investigation: J.S., Y.-J.P., M.A.T., L.Z., S.M., A.Addetia, V.V.P., C.A.T., J.B., C.S., S.R.L, B.M., A.Aka, A.M.J.

Validation: J.S., Y.-J.P., M.A.T., D.V., S.L.

Resources: A.Aka, Z.Z., E.F.F., A.M.J., C.B.W., D.V., S.L.

Funding acquisition: D.V., S.L. Supervision: D.V., S.L.

Writing – original draft: J.S., Y.-J.P., D.V., S.L.

Writing – review & editing: J.S., Y.-J.P., M.A.T., C.B.W., D.V., S.L.

